# Towards a combined therapy for spinal muscular atrophy based on opposing effects of an antisense oligonucleotide on chromatin and splicing

**DOI:** 10.1101/2021.09.24.461646

**Authors:** Luciano E. Marasco, Gwendal Dujardin, Rui Sousa-Luís, Ying Hsiu Liu, José Stigliano, Tomoki Nomakuchi, Nick J. Proudfoot, Adrian R. Krainer, Alberto R. Kornblihtt

**Affiliations:** Universidad de Buenos Aires (UBA), Facultad de Ciencias Exactas y Naturales, Departamento de Fisiología, Biología Molecular y Celular and CONICET-UBA, Instituto de Fisiología, Biología Molecular y Neurociencias (IFIBYNE), Buenos Aires, Argentina; Sir William Dunn School of Pathology, Oxford, UK; Instituto de Medicina Molecular, Universidade de Lisboa, Portugal; Cold Spring Harbor Laboratory, Cold Spring Harbor, New York, USA

## Abstract

Spinal Muscular Atrophy (SMA) is a motor-neuron disease caused by loss-of-function mutations of the *SMN1* gene. Humans have a paralog, *SMN2*, whose exon 7 is predominantly skipped, and so it cannot fully compensate for the lack of *SMN1*. Nusinersen (Spinraza) is a splicing-correcting antisense oligonucleotide drug (ASO) approved for clinical use. Nusinersen targets a splicing silencer located in *SMN2* intron 7 pre-mRNA and, by blocking the binding of the splicing repressors hnRNPA1 and A2, it promotes higher E7 inclusion, increasing SMN protein levels. We show here that, by promoting transcriptional elongation, histone deacetylase (HDAC) inhibitors cooperate with a nusinersen-like ASO to upregulate E7 inclusion. Surprisingly, the ASO also elicits the deployment of the silencing histone mark H3K9me2 on the *SMN2* gene, creating a roadblock to RNA polymerase II elongation that acts negatively on E7 inclusion. By removing the roadblock, HDAC inhibition counteracts the undesired chromatin effects of the ASO, resulting in higher E7 inclusion. Combined systemic administration of the nusinersen-like ASO and HDAC inhibitors in neonate SMA mice had strong synergistic effects on SMN expression, growth, survival, and neuromuscular function. Thus, we suggest that HDAC inhibitors have the potential to increase the clinical efficacy of nusinersen, and perhaps other splicing-modulatory ASO drugs, without large pleiotropic effects, as assessed by genome-wide analyses.

## Introduction

Spinal Muscular Atrophy (SMA) is a motor-neuron disease caused by loss-of-function mutations of the *SMN1* gene. Humans have a paralog, *SMN2*, whose exon 7 (E7) is predominantly skipped, and so it cannot fully compensate for the lack of *SMN1*. Both SMN2 and SMN1 are about 30 kbp-long each but they differ in only 11 bp, which indicates a recent duplication in evolution. The only critical difference between the two *SMN* genes is the C>T base change 6 bp inside exon 7 that causes E7 skipping (Monani et al., 2000), giving rise to a truncated and inactive version of the SMN protein. Both genes encode a splicing silencer sequence in intron 7 that is the target site for the negative splicing factors hnRNPA1 and A2 that contribute to E7 skipping in *SMN2*. This basic mechanistic knowledge led to the development of an antisense oligonucleotide (ASO) therapeutic strategy for SMA: Nusinersen (Spinraza) is a splicing-correcting ASO drug approved for clinical use. Nusinersen targets a splicing silencer located in *SMN2* intron 7 pre-mRNA and, by blocking the binding of hnRNPA1 and A2, it promotes higher E7 inclusion, increasing SMN protein levels. Nusinersen was the first drug approved for SMA therapy and also the first-splicing-corrective drug. Although nusinersen is administered to patients by lumbar puncture, to readily reach motor neurons, systemic administration robustly rescues severe symptoms in an SMA mouse model (Hua et al., 2008, 2011, 2015) which suggests that restoring adequate SMN levels in peripheral tissues might be therapeutically important.

Alternative splicing is not only regulated by the binding of activators and repressors to splicing enhancers and silencers, but also by the rate of RNA polymerase II (RNAPII) transcriptional elongation and chromatin structure. According to the kinetic coupling mechanism, slow elongation can promote either exon inclusion or skipping, depending on the type of exon (de la Mata et al., 2003; Dujardin et al., 2014, Fong et al., 2014). In class I exons, slow elongation promotes inclusion by improving the recruitment of constitutive or regulatory splicing factors to the nascent mRNA, whereas in class II exons slow elongation enhances the binding of inclusion inhibitory factors to their target sites. Genome-wide studies showed that approximately 20% of a cell’s alternative splicing events are sensitive to elongation and, depending on the cell type, class I exons represent between 50 and 80%, while class II exons are between 20 to 50% of the elongation-sensitive alternative exons (Fong et al, 2014; Ip et al., 2011, Muñoz et al. 2009, 2017; Maslon et al., 2019). On the other hand, intragenic deployment of specific histone marks can create more compact or more relaxed chromatin regions that inhibit or promote RNAPII elongation, respectively and consequently may affect alternative splicing. For example, acetylation of histone H3 lysine 9 (H3K9Ac) along gene bodies promotes skipping of class I exons (Schor et al., 2009), whereas dimethylation of the same residue (H3K9me2) promotes their inclusion (Alló et al., 2009; Schor et al., 2013). Conversely, intragenic H3K9 acetylation promotes inclusion of class II exons, which is consistent with the fact that treatments that inhibit elongation promote their skipping (Dujardin et al., 2014).

We show here that *SMN2* E7 is a class II alternative exon, which opened the way to explore the regulation of its inclusion using kinetic coupling tools. We found that, by promoting transcriptional elongation, histone deacetylase (HDAC) inhibitors cooperate with a nusinersen-like ASO to upregulate E7 inclusion. Surprisingly, the ASO also elicits the deployment of the silencing histone mark H3K9me2 on the *SMN2* gene, creating a roadblock to RNA polymerase II elongation that acts negatively on E7 inclusion. By removing the roadblock, HDAC inhibition counteracts the undesired chromatin effects of the ASO, resulting in higher E7 inclusion. Combined systemic administration of the nusinersen-like ASO and HDAC inhibitors in neonate SMA mice had strong synergistic effects on SMN expression, growth, survival, and neuromuscular function. Thus, we suggest that HDAC inhibitors have the potential to increase the clinical efficacy of nusinersen, and perhaps other splicing-modulatory ASO drugs, without large pleiotropic effects, as assessed by genome-wide analyses.

## Results and Discussion

### Slow elongation inhibits *SMN2* E7 inclusion

To evaluate the impact of elongation rate on *SMN2* E7 inclusion, we assessed the effects of a slow mutant of RNAPII. We co-transfected HEK293T cells with an *SMN2* E7 reporter minigene (Lorson et al. 1999) and a plasmid encoding an α-amanitin-resistant large subunit of RNAPII, with or without the point mutation R749H (de la Mata et al., 2003), previously shown to slow elongation in vivo by 2- to 3-fold (Boireau et al., 2007; Maslon et al., 2019). When transcription was carried out by the slow polymerase, *SMN2* E7 skipping was greater than upon transcription by the α - amanitin-resistant wild-type polymerase (Fig. 1A). The use of a minigene splicing reporter is necessary to assess the effects of the slow polymerase by transient co-transfections. However, and more importantly, the effect is also observed on the endogenous *SMN2* E7 as shown by re-analysis of published genome-wide sequencing data (Fong et al., 2014) that confirmed that not only the slow polymerase causes E7 skipping, but that a fast mutant (E1126G) has the expected opposite effect, promoting E7 inclusion (Fig. 1B). Consistently, treatment of cells with the DNA topoisomerase I inhibitor camptothecin (CPT), which indirectly inhibits elongation (Listerman et al., 2006; Dujardin et al. 2014), also promoted E7 skipping from the endogenous *SMN2* gene (Fig. 1C). These results indicate that *SMN2* E7 is a class II exon.

**Figure 1.**
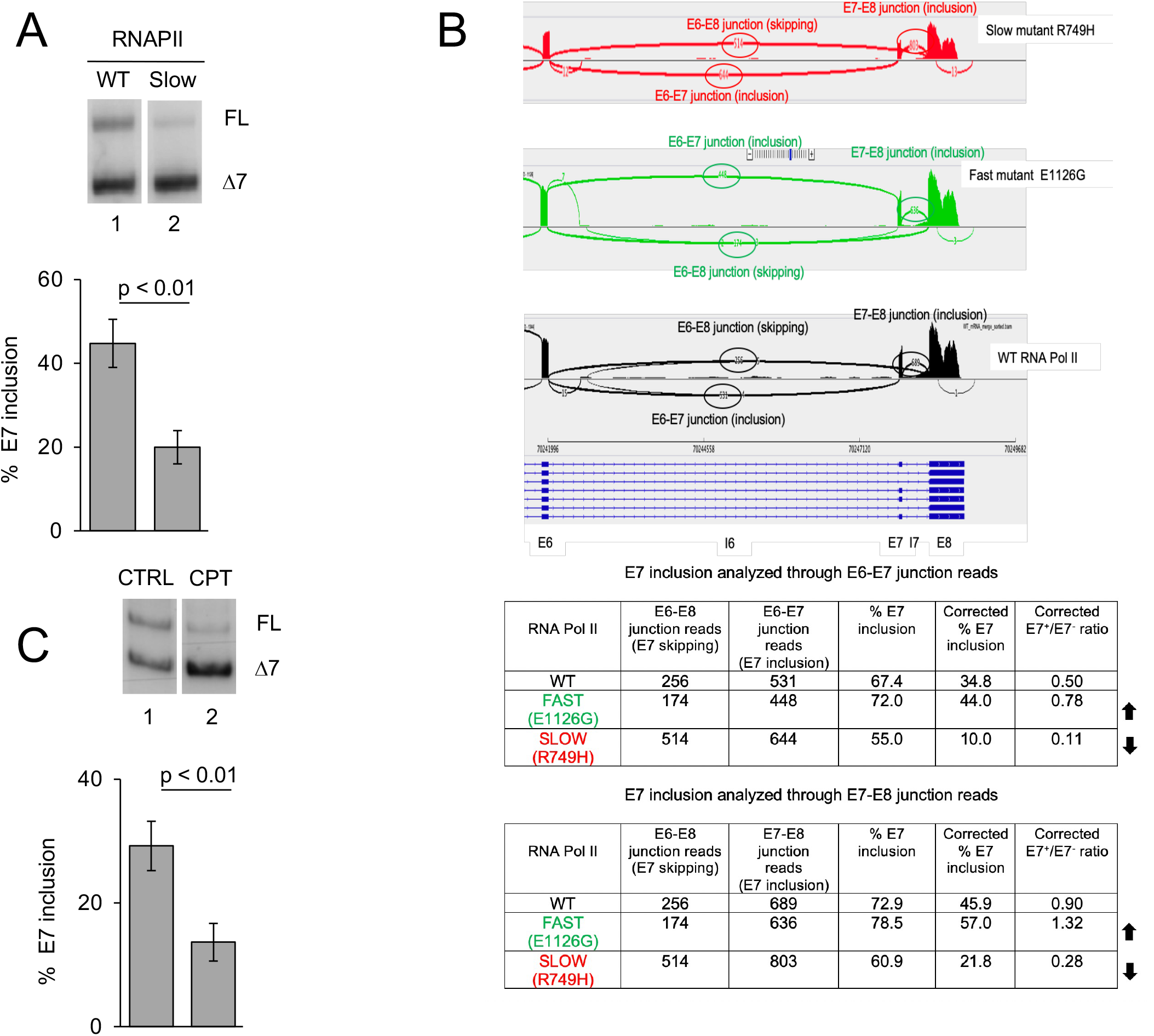
*SMN2* E7 inclusion into the mature mRNA is controlled by RNAPII elongation behaving as a class II exon. (A) Transcription by a slow RNAPII mutant inhibits *SMN2* E7 inclusion. HEK293T cells were co-transfected with the *SMN2* reporter minigene and expression vectors for WT^res^ and hC4 RNAPII (slow mutant), followed by addition of α-amanitin. Alternative splicing was analyzed by radioactive RT-PCR followed by native polyacrylamide gel electrophoresis and autoradiography. Bars display means ± SD of percentage of the radioactivity in the full-length (FL) band, containing E7, over the sum of radioactivity in the FL and ΔE7 (lacking E7) bands of at least three independent transfection experiments. A two-tailed Student’s t test was used to determine the significance between events. (B) Re-analysis of the RNA-seq data published by the Bentley laboratory (Fong et al., 2014). Due to the almost identical DNA sequences of the human *SMN1* and *SMN2* genes (only 11 bp differences in ∼30 kbp), sequencing reads were assigned to a merge of the two genes. For this reason, splicing junction data had to be corrected considering 100% inclusion in the case of the *SMN1* gene. Top: “Sashimi plots” indicating the number of reads for the E6-8 (skipping), E7-E8 (inclusion) and E6-E7 (inclusion) junctions of the *SMN*1/2 merged gene in cells stably transfected with the slow (R749H, in red), fast (E1126G, in green) and WT (in black) RNAPII large subunit constructs, all of which had a second mutation that confers resistance to α-amanitin, a drug that was used to treat the cells before RNA extraction. Bottom: Raw and corrected quantification of the levels of E7 inclusion as assessed by analysis of the E6-E7 junction reads (top) and E7-E8 junction reads (bottom). Inclusion levels are expressed as both percentage of E7 inclusion and E7^+^ / E7^-^ (i.e. FL/ Δ7) ratios. (C) Effect of camptothecin (CPT)on alternative splicing of endogenous *SMN2* E7. HEK293T cells were incubated for at least 6 hours with 3 µM CPT. Alternative splicing was assessed as in (A).

### Cooperative effects of ASO1 and HDAC inhibitors

In view of the above results, we reasoned that chromatin opening by histone acetylation should foster E7 inclusion by promoting RNAPII elongation. Therefore, we measured the effects of HDAC inhibitors, such as trichostatin A (TSA), on E7 inclusion in HEK293T cells, either alone or in combination with transfection of a nusinersen-like 2’-O-(2-methoxyethyl) (MOE) phosphorothioate-modified antisense oligonucleotide (named ASO1 in this paper) at suboptimal concentrations. Nusinersen (5’-TCACTTTCATAATGCTGG-3^′^) was originally dubbed ASO10-27 (Hua et al., 2008), and the equally effective ASO1 variant (5^′^-ATTCACTTTCATAATGCTGG-3^′^) has a 2-nt extension and can be described as ASO10-29. TSA alone was less potent than ASO1 in upregulating E7 inclusion, but combining both drugs greatly enhanced the effect (Fig. 2A). The cooperative effect of TSA was dose-dependent (Suppl. Fig. 1A) and reproducible in other cell types, such as HeLa (Suppl. Fig. 1B) and SMA patient fibroblasts (Suppl. Fig. 1C). We obtained similar results by combining ASO1 and other HDAC inhibitors such as SAHA (N-hydroxy-N•-phenyl-octanediamide) (not shown) and valproic acid (VPA, Fig. 2B). Unlike TSA and SAHA, VPA is approved for clinical use (Wirth et al., 2006). Also, similar effects were obtained with an ASO with the same sequence as ASO1 but with a different chemical backbone (2’-O-methyl phosphorothionate) (not shown).

**Figure 2.**
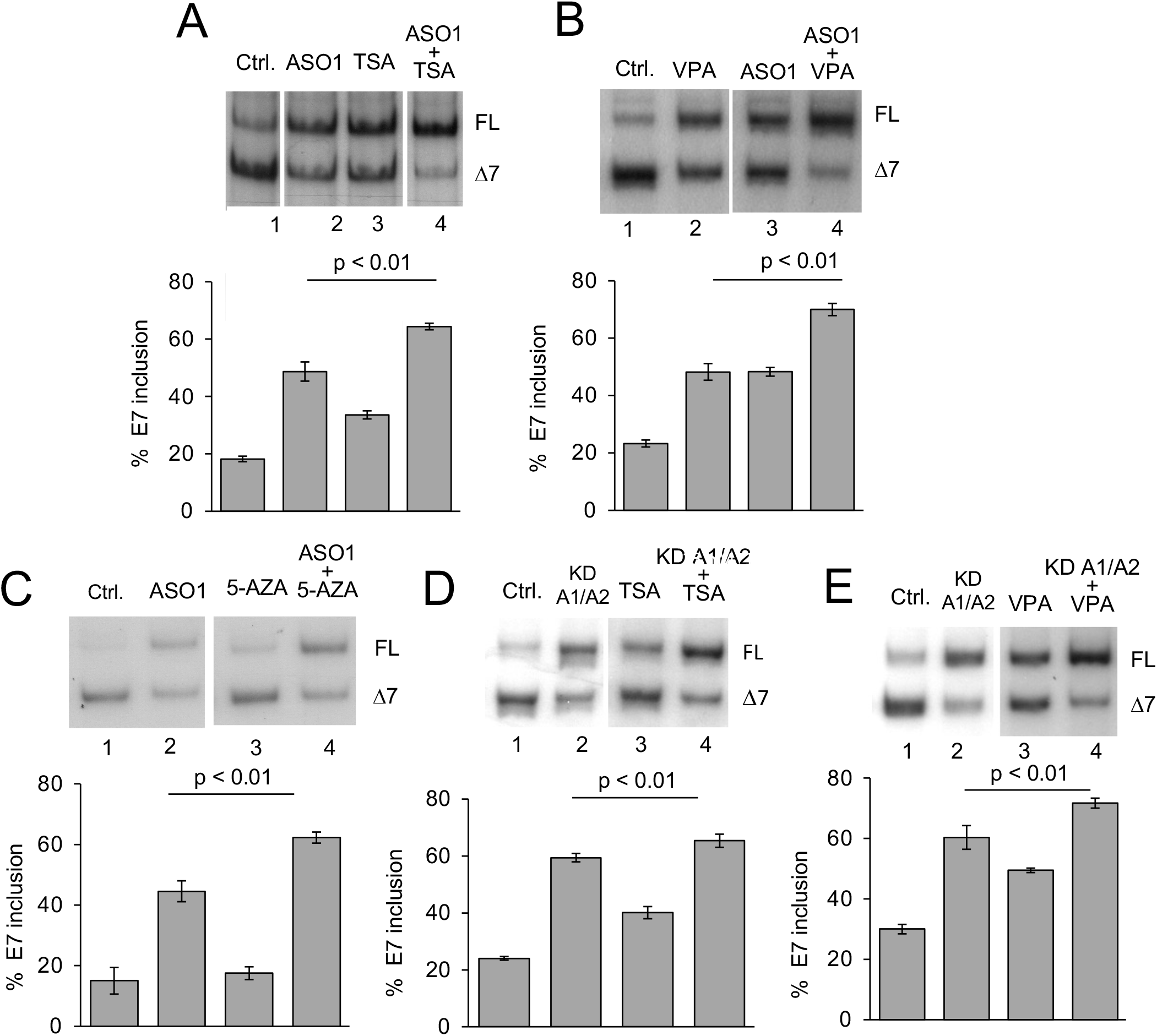
HDAC and histone methylation inhibitors potentiate ASO1’s upregulation of E7 inclusion. Combined effects on endogenous *SMN2* E7 alternative splicing of transfection with 25 nM ASO1 and treatment with 3 µM TSA for 24 hr (A), 10 mM VPA for 24 hr (B), or 25 µM 5-AZA for 48 hr (E). Combined effects on HEK293T endogenous *SMN2* E7 alternative splicing of transfection with 25 nM each of si-hnRNPA1 and si-hnRNPA2, and treatment with 3 µM TSA for 24 hr (F) or 10 mM VPA for 24 hr (G). An oligonucleotide with a ASO1 scrambled sequence was used as control (scrambled oligo). Alternative splicing was assessed as in Fig. 1A.

H3K9me2 is a transcriptionally repressive mark that competes with the transcriptionally permissive H3K9Ac (Ghare et al., 2014). Accordingly, with the caveat of potential indirect effects, inhibition of methylation by 5-aza cytidine (5-AZA), a drug that inhibits both DNA and histone methylation (Wozniak et al., 2007) enhanced the effects of ASO1 as effectively as the HDAC inhibitors (Fig. 2C). Finally, in agreement with the fact that ASO1 displaces the splicing repressors hnRNPA1/A2 from their pre-mRNA target site in intron 7, we found that upon treatment with either TSA (Fig. 2D) or VPA (Fig. 2E), the effects of hnRNPA1/A2 depletion in promoting E7 inclusion were stronger.

VPA was previously assessed in the context of SMA, but not in combination with nusinersen. Clinical trials with VPA failed to demonstrate effectiveness (Swoboda et al., 2010; Kissel et al., 2011). The rationale for VPA’s clinical use was that chromatin opening at the *SMN2* promoter would increase transcription and therefore SMN levels (Kernochan et al., 2011). Both TSA and VPA indeed increased the levels of *SMN2* pre-mRNA in HEK293T cells, measured with a promoter-proximal amplicon as a proxy to transcription levels (Suppl. Fig. 1 D and E). However, it should be noted that HDAC inhibitors alone do not seem to have therapeutic effects in SMA mice (see below). ChIP analysis (Fig. 3A) shows that VPA promoted intragenic H3K9 acetylation along the *SMN1/2* gene, with a conspicuous peak around the E7 region, which may explain why E7 inclusion is upregulated, according to the kinetic-coupling mechanism of alternative splicing (Kornblihtt et al., 2013). We ruled out the possibility that the HDAC inhibitors may be acting through modulation of SMN protein levels, which in turn might affect *SMN2* E7 splicing (Jodelka et al. 2010), because overexpressing SMN did not alter the effect of ASO1 (Suppl. Fig. 2A)

**Figure 3.**
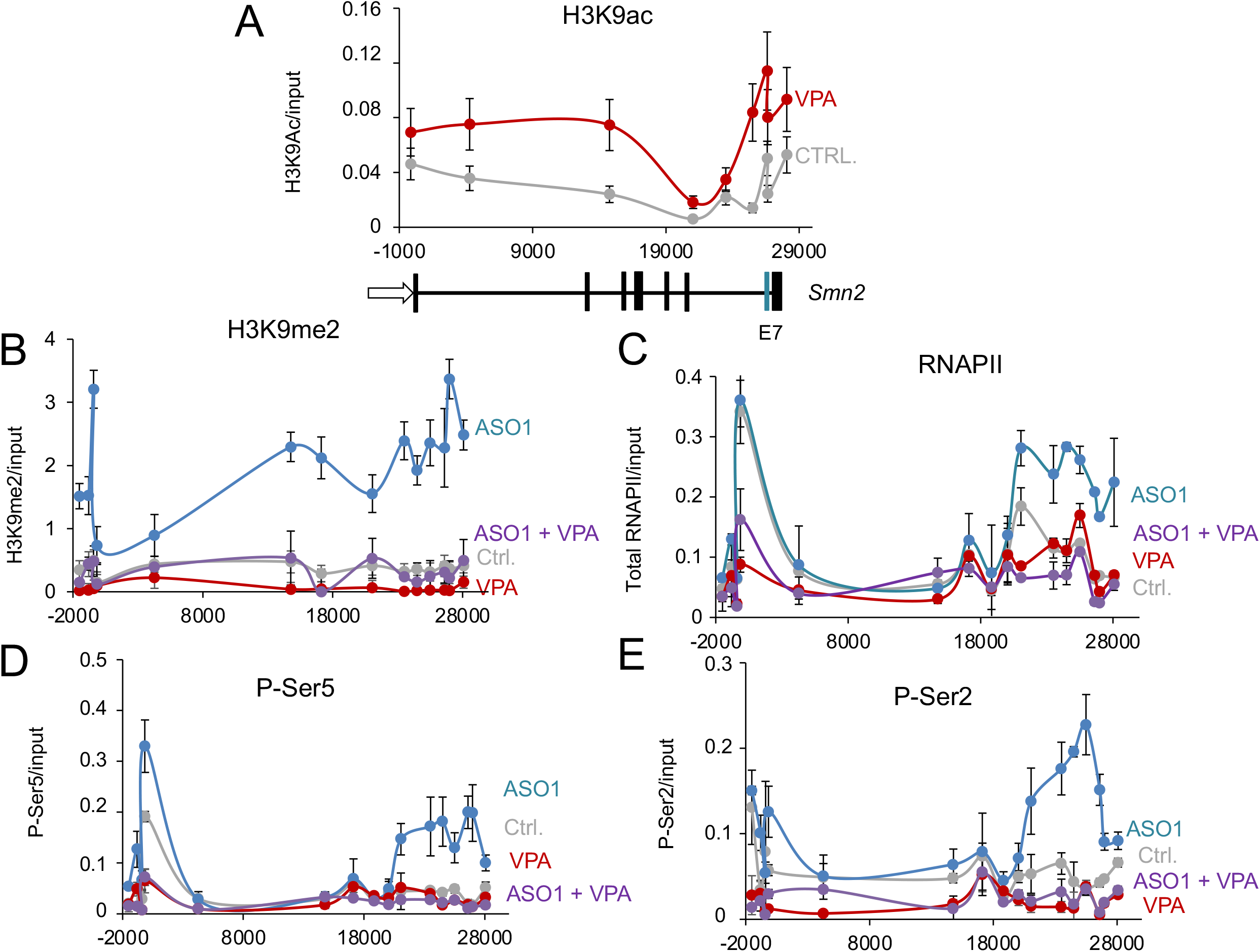
ASO1 promotes H3K9 dimethylation along the *SMN1/2* gene, creating a roadblock to RNAPII. (A) H3K9Ac distribution along the *SMN2* gene, assessed by ChIP-qPCR in HEK293T cells treated (red) or untreated (gray) with 10 mM VPA for 12 hr. H3K9me2 (B), total RNAPII (C), P-Ser5 RNAPII (D), and P-Ser2 RNAPII (E) distribution along the *SMN2* gene, assessed by ChIP-qPCR in HEK293T cells transfected with the scrambled oligo (CTRL, gray) or ASO1 (blue and purple) and treated with 10 mM VPA (red and purple). Four independent immunoprecipitation replicates were conducted per experiment. Data are represented as mean ± S.D. (n = 4, *p < 0.05, two-tailed Student’s t test).

### Chromatin effects of ASO1

Due to the fact that ASO1 is a single-stranded oligonucleotide that hybridizes with an intronic sequence at the pre-mRNA level, we wondered if it could have chromatin effects similar to those of siRNAs directed to intronic regions, as previously described (Alló et al., 2009, Schor et al., 2013). These reports showed that siRNAs whose guide strands are complementary to pre-mRNA intronic sequences (antisense) located downstream of an alternative exon regulate the splicing of that exon by promoting silencing marks, such as H3K9me2, which in turn act as roadblocks to RNAPII elongation. This effect was shown to be counterbalanced by factors that favor chromatin opening or transcriptional elongation. Remarkably, here we observed that transfection of HEK293T cells with ASO1 promoted extensive H3K9 dimethylation along the *SMN1/2* gene, with peaks that reached an 8-fold increase at the promoter and the alternative E7 areas (Fig. 3B, cf. blue and gray lines). Other histone marks, like H3K27me3 or H3K9Ac, were not affected by ASO1 transfection (Suppl. Fig. 2B). Most importantly, VPA not only reduced the H3K9me2 marks relative to the control (Fig. 3B, cf. red and gray lines) but completely abolished the effect of ASO1 (Fig. 3B, cf. blue and purple lines), confirming antagonistic roles of histone methylation and acetylation. These findings reveal unforeseen alterations to the deployment of histone marks at the targeted gene, induced by ASO treatment. Notably these effects that can be abrogated by chromatin opening with HDAC inhibitors.

We next tested whether RNAPII densities were altered. Transfection with ASO1 greatly increased total, P-Ser5 and P-Ser2 (Figs. 3C-E) RNAPII densities at the promoter and at a distinct peak near the ASO1 target site on the transcript. In all three cases, the RNAPII peak was abolished when cells were additionally treated with VPA. We interpret that by promoting H3K9 methylation, ASO1 creates a roadblock to RNAPII upstream of its target site. This may equally affect the total and the two main phosphoisoforms of RNAPII, suggesting that it acts as a general steric impediment, rather than regulating RNAPII phosphorylation. The increase in H3K9me2 and RNAPII at the promoter region, which does not have a target site of ASO1, may reflect looping between the two ends of the gene (Grzechnik et al., 2014). RNase H, an enzyme that degrades RNA in RNA-DNA hybrids, did not alter the effect of ASO1 on E7 inclusion (Suppl. Fig. 2C), which suggests that the chromatin effect of ASO1 does not appear to involve R-loop formation (Tang-Wong et al., 2019). This might indicate that ASO1 does not act through hybridization to DNA, as opposed to RNA, consistent with the previous demonstration that an oligonucleotide with complementary sequence to nusinersen (sense oligo) has no effect on *SMN2* E7 inclusion (Rigo et al., 2014). Moreover, the effect of ASO1 was unchanged by depletion of AGO1 (Suppl. Fig. 2D), known to be required for the chromatin effects of siRNA (Alló et al., 2009).

### Uncoupling the two opposite roles of ASO1

The above results suggest a model whereby in the absence of HDAC inhibitors, ASO1 has two opposing effects on E7 inclusion (Fig. 4A, left panel). It promotes inclusion by blocking hnRNPA1 and A2 binding to the pre-mRNA, but concomitantly results in a compact chromatin structure and a roadblock to elongation that in turn promotes E7 skipping. Although both effects co-exist, at high ASO1 concentrations, the chromatin effect is evidently surpassed by the hnRNPA1/A2 effect. On the other hand, in the presence of HDAC inhibitors (right panel), the chromatin-silencing effect is abrogated, and the levels of E7 inclusion increase further. To test the validity of this model, we investigated the effects of a second ASO, ASO2, whose target site is also located in intron 7, but downstream of the ASO1 target site (Suppl. Fig. 3A). ASO2 bears no sequence identity with ASO1, and does not overlap hnRNPA1/A2 binding sites. Even so, if ASO2 elicits the chromatin effect, it should inhibit E7 inclusion. Fig. 4B shows that, whereas ASO1 increases, ASO2 inhibits E7 inclusion. Furthermore, similarly to ASO1, ASO2 promoted H3K9 methylation (Fig. 4C) and higher total, P-Ser5, and P-Ser2 (Figs. 4D-F) RNAPII densities, consistent with roadblocks to elongation that in all cases were abolished by treatment with VPA. As specificity controls, neither ASO1 nor ASO2 promoted H3K9 dimethylation in a gene located in the same topologically-associated domain (TAD) as *SMN2* (Lefebvre et al., 1995) (Suppl. Fig. 3B), or in genes located outside the *SMN2* TAD with either high (Suppl. Fig. 3C) or low (Suppl. Fig. 3D) basal levels of H3K9 methylation.

**Figure 4.**
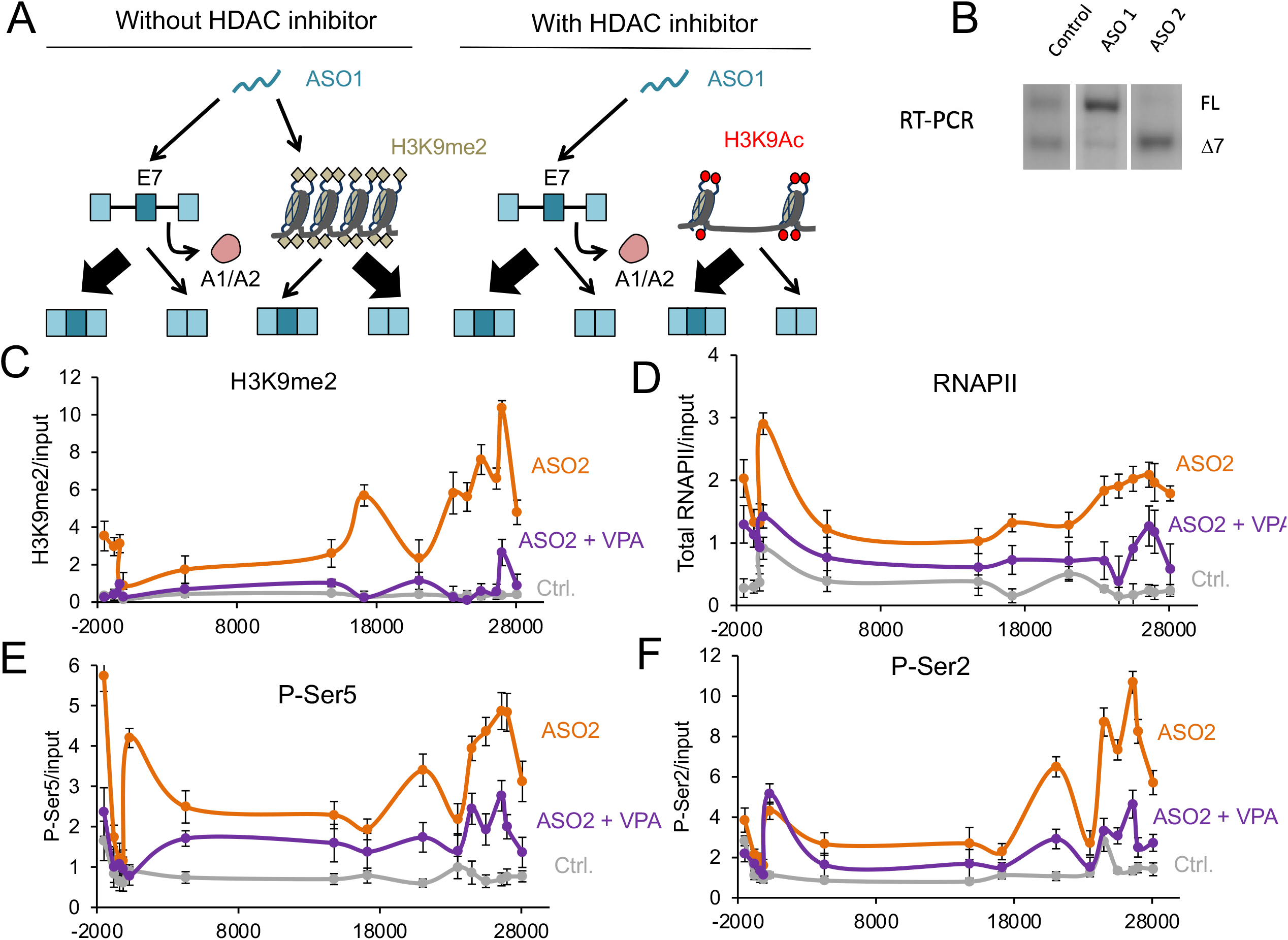
Uncoupling the two opposing effects of ASO1: ASO2 also promotes H3K9 dimethylation and imposes an RNAPII roadblock, but inhibits *SMN2* E7 inclusion. (A) Model of two opposing roles of ASO1. In the absence of HDAC inhibition, ASO1 promotes chromatin condensation (left); in the presence of HDAC inhibitors, chromatin becomes more relaxed, counteracting the condensation promoted by ASO1 (right). (B) Effects on endogenous *SMN2* E7 alternative splicing of transfecting HEK293T cells with with 25 nM ASO2 using the scrambled oligo and ASO1 as negative and positive controls respectively. Alternative splicing was assessed like in Fig. 1A. H3K9me2 (C), total RNAPII (D), P-Ser5 RNAPII (E), and P-Ser2 RNAPII (F) distribution along the *SMN1/2* gene, assessed by ChIP-qPCR in HEK293T cells transfected with the scramble oligo (CTRL, gray) or ASO2 (orange and purple) and treated with 10 mM VPA (red and purple). Four independent immunoprecipitation replicates were conducted per experiment. Data are represented as mean ± S.D. (n = 4, *p < 0.05, two-tailed Student’s t test).

These uncoupling experiments with ASO2 not only confirm the model in Fig. 4A, but further highlight the importance of potential chromatin effects when evaluating antisense therapeutic strategies.

To confirm the mechanism proposed in Fig. 4A, we next performed CRISPR ablation of the gene encoding the histone H3K9 dimethyl transferase G9a (Fiszbein et al., 2016). The effectiveness of this gene knock out was confirmed by western blot (Fig. 5A). Notably this promoted the upregulation of *SMN2* E7 inclusion by ASO1 (Fig. 5B). When plotted as the E7+/E7-ratio (Fig. 5B, bottom), the absence of G9a increased E7 inclusion by 4-fold.

**Figure 5.**
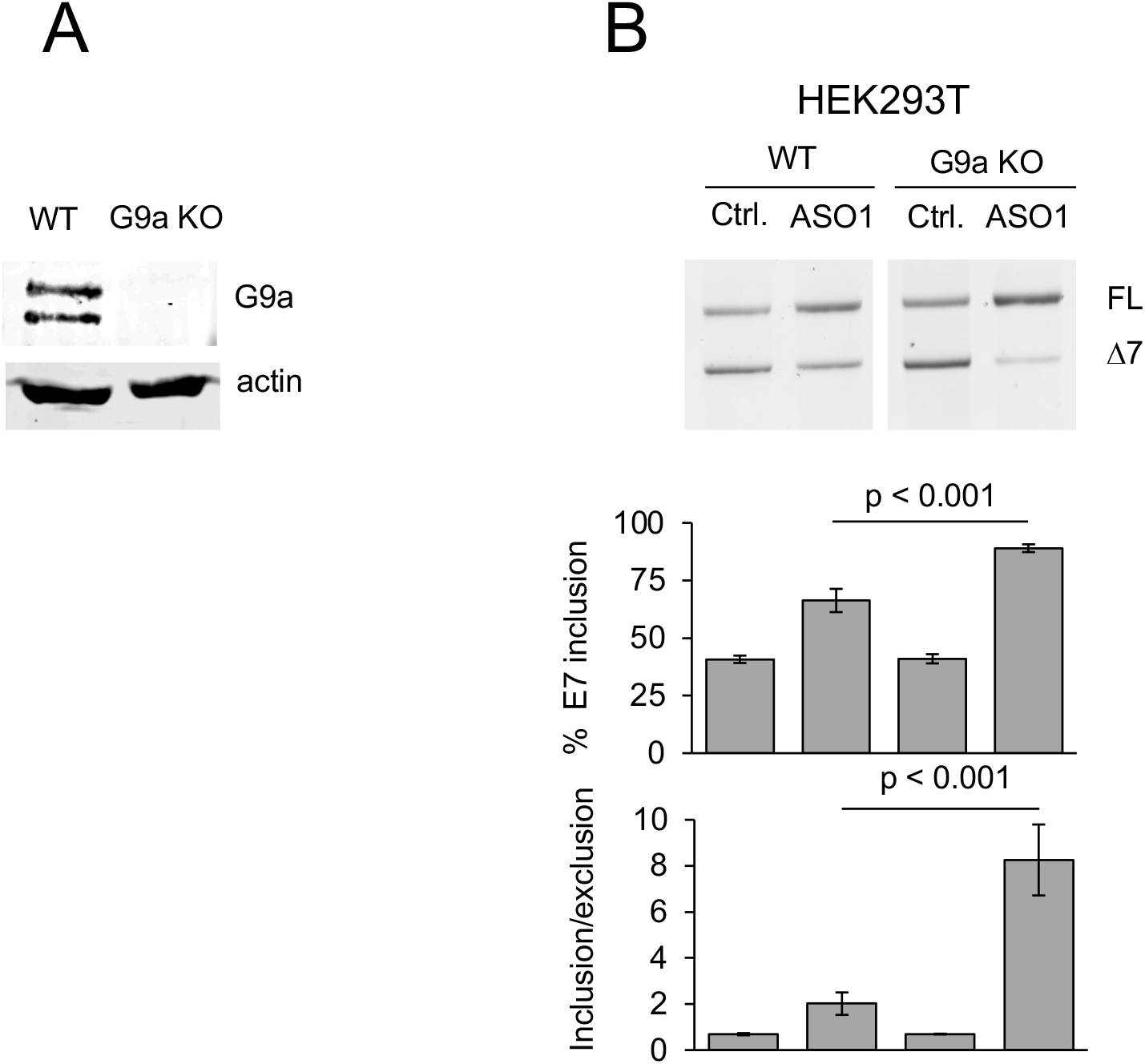
Ablation of the H3K9 dimethyl transferase G9a potentiates upregulation of *SMN2* E7 inclusion by ASO1. (A) Western blot showing the absence of G9a protein in the cells in which the G9a gene was knocked out. (B) Alternative splicing was analyzed, as in Fig. 1A, in WT and CRISPR G9a-ablated HEK293T cells using the scrambled oligo as control. Bars display means ± SD of the percentage of the radioactivity in the FL band over the sum of radioactivity in the FL and Δ7 bands (top) or as E7 inclusion over exclusion ratios (bottom) of at least three independent transfection experiments.

### Genome-wide analyses

We performed genome-wide characterization of the effects of ASO1 and VPA in HEK293T cells, including nascent-mRNA profiles, mRNA expression, and alternative splicing, using the recently developed POINT-seq technology (Sousa-Luís et al., 2021) as well as more standard RNA-seq. Metagene analysis of POINT-seq on 2741 expressed protein-coding genes confirmed that VPA dramatically changed the pre-mRNA read-count ratio between the promoter region (P) and the gene body (GB) (Fig. 6A), whereas ASO1 did not (Fig. 6B). Similar results were obtained when assessing the number of genes with different P/GB values for control versus VPA- or ASO1-treated cells (Fig. 6C).

**Figure 6.**
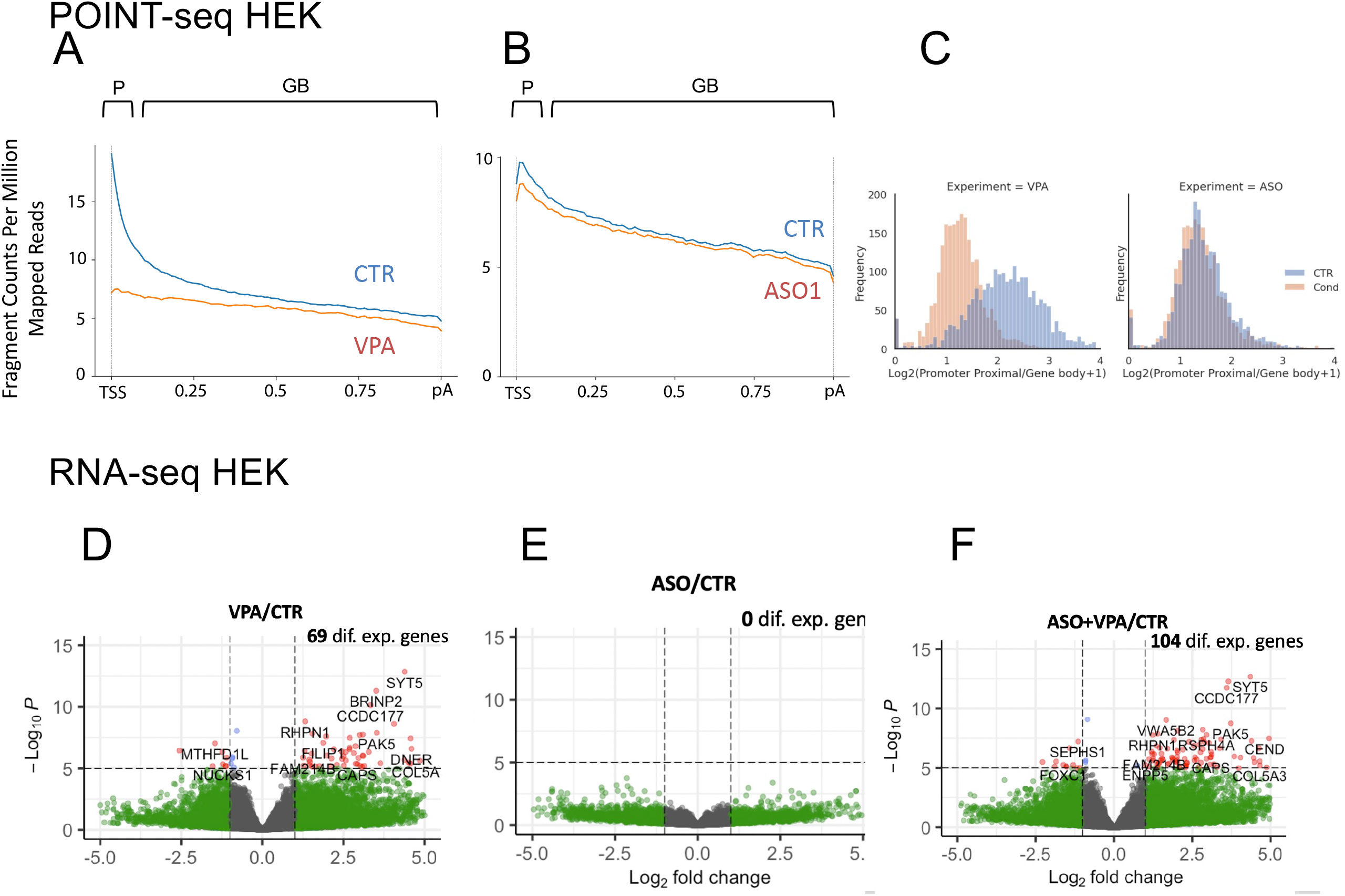
Global effects of VPA and ASO1 on transcription elongation and gene expression. Metagene analysis of POINT-seq signal in normalized transcription units from the transcription start site (TSS) to the cleavage and polyadenylation site (pA) of non-overlapping protein-coding genes of HEK293T cells treated or not with VPA (A) or ASO1 (B). (C) Histograms representing the log2 ratio distribution of read density between the promoter-proximal region (P) and gene body (GB) for POINT-seq samples. VPA experiment is shown on the left, and ASO1-treated cells on the right. Invalid log2 values were avoided by adding 1 unit to each ratio. (D-F) RNA-seq volcano plots showing differentially expressed genes in HEK293T cells treated with VPA (D), ASO1 (E) and VPA + ASO1 (F) with respect to control untreated cells. Only red points represent significantly differentially expressed genes, by showing a p-value lower than 1 × 10^−5^ and an absolute log2FoldChange value higher than 1. Green points represent genes for which a fluctuation between conditions was observed, but it was inconsistent between replicates, either due to technical variability or a low number of reads. Grey dots are genes with log2FoldChange smaller than 1. Three replicates per condition were employed.

RNA-seq analysis revealed that only modest changes in gene expression and alternative splicing were detected. Thus only 69 genes out of 29,000 transcription units analyzed were altered by VPA treatment (Fig. 6D). It is worth mentioning here that the computing tools allowing to assess gene expression by RNA-seq analysis exclude by default reads mapping to duplicated genome sequences. For this reason, we will not see here the mild increase in *SMN1/2* expression caused by HDAC inhibitors, revealed by RT-qPCR in Suppl. Figs 1D and E. Notably none of the 29,000 transcription units significantly changed expression in response to ASO1 (Fig. 6E), and the combined effects of VPA + ASO1 were also minimal (104 genes over 29,000 transcription units, Fig. 6F). In this case there is full consistency with the individual analysis of *SMN1/2* gene because ASO1 had no effect (Suppl. Figs 1D and E). One may wonder why there are so few genes changing expression. We think that although VPA affects elongation, there is no evidence that it promotes higher RNAPII recruitment, which may explain why we previously saw that faster elongation affected the quality (splicing isoform balance) but not the quantity of NCAM mRNA (Schor et al., 2009). In any case, the concentrations and short times of our VPA treatment are sufficient to enhance the effect of ASO1 on splicing without major expression alterations. In fact, comparable RNA-seq results were obtained in other studies, under similar VPA treatments. For example, Balasubramanian et al. (2019) reported that treatment of RN46A cells with 0.5 mM VPA for 72 h caused regulation of only 88 of the ∼16,500 genes analyzed. However, treatments of rats with VPA concentrations 60 times higher than the ones we used in mice here (see below) change expression of about 3,000 genes (Zhang et al., 2018) which suggests a concentration-dependent outcome at the gene regulation level.

Similarly, RNA-seq revealed that VPA only affected the patterns of 48 of the 79,549 alternative splicing events analyzed (0.06%, Suppl. Figs. 4A and B), when a relatively high threshold (□PSI>30%) is applied, and confirmed the cooperative effects of ASO1 and VPA on *SMN2* E7 inclusion (Suppl. Figs. 4C and D). In conclusion, these data are consistent with a global increase in transcription elongation caused by VPA, as previously shown for individual genes, where elongation speed increased from ∼2 to ∼4 kb/min the with HDAC inhibitor TSA (Dujardin et al., 2014), but with no conspicuous global changes in gene expression and alternative splicing. In parallel, transfection with ASO1 had no global effects on elongation or gene expression.

The recently designed POINT-seq method assesses the distribution of nascent mRNA reads associated to elongating RNAPII along transcribed genes. Since the *SMN1* and *SMN2* genes only differ in 11 bp in their similar lengths of about 30,000 bp each, it becomes impossible to assign POINT-seq reads individually to each of the two genes. For these reasons, POINT-seq analysis must necessarily refer to the *SMN1/2* merged genes. In spite of these limitations, POINT-seq data showed that, unlike ASO1, VPA reduced the ratio of read counts at the Promoter over the Gene Body (P/GB ratio) from 4.17 to 1.73 (Suppl. Fig. 5A), consistent with the global effect of VPA on elongation. However, when focusing on the region around ASO1’s target site (Suppl. Fig. 5B), we clearly observe that ASO1 increased the ratio of reads mapping upstream versus downstream of the target site (U/D ratio), which is highly consistent with the roadblock to elongation evidenced by RNAPII ChIP (Fig. 3).

Overall, our genome-wide analyses reinforce the evidence that the combined treatment with ASO1 and HDAC inhibitors enhances *SMN2* E7 inclusion, accompanied by only marginal alterations—both in the number of genes and in the magnitude of the change—of global gene expression and alternative splicing.

### Combined treatment in SMA mice

Nusinersen, administered subcutaneously in newborn SMA mice, distributes broadly and strongly rescues survival and motor function (Hua et al., 2011). We therefore sought to test the combined effect of ASO1 and VPA treatment using a mouse model of severe SMA. Mice have only one *Smn* gene, and *Smn*^*-/-*^ mutants are embryonic lethal (Hsie-Li et al., 2000; Monani et al., 2000). We used *Smn*-null transgenic mice containing two copies of the human *SMN2* transgene, which develop a severe SMA-like phenotype with a mean survival of ∼7-10 days (Hua et al., 2011). We administered two consecutive low-dose subcutaneous injections of ASO1 (18 µg/g) at P0 and P1 and/or one subcutaneous injection of VPA (10 µg/g) at P1. The rationale for the choice of a low or suboptimal dose of ASO1 is to elicit a partial phenotypic effect that allows us to evaluate improvements by co-administration of HDAC inhibitors. Since the generation of mice with the two mouse *SMN* alleles disrupted is performed by crossing of *Smn*^*+/-*^ heterozygous and the *Smn*^*-/-*^ (SMA mice) and *Smn*^*-/+*^ phenotypes are not distinguished at birth, injections were blind with respect to the genotypes and genotyping was performed after injections. Fig. 7A shows that SMA mice injected with VPA alone died before P10, similarly to vehicle-treated controls. The suboptimal dose of ASO1 alone extended median survival to ∼20 days, while co-injection of VPA with ASO1 increased the median survival to ∼70 days. All ASO1-alone-injected mice were dead at P62, whereas ∼60% of those treated with ASO1 and VPA were still alive at P62. The difference between the two treatments is statistically significant (p = 0.00081).

**Figure 7.**
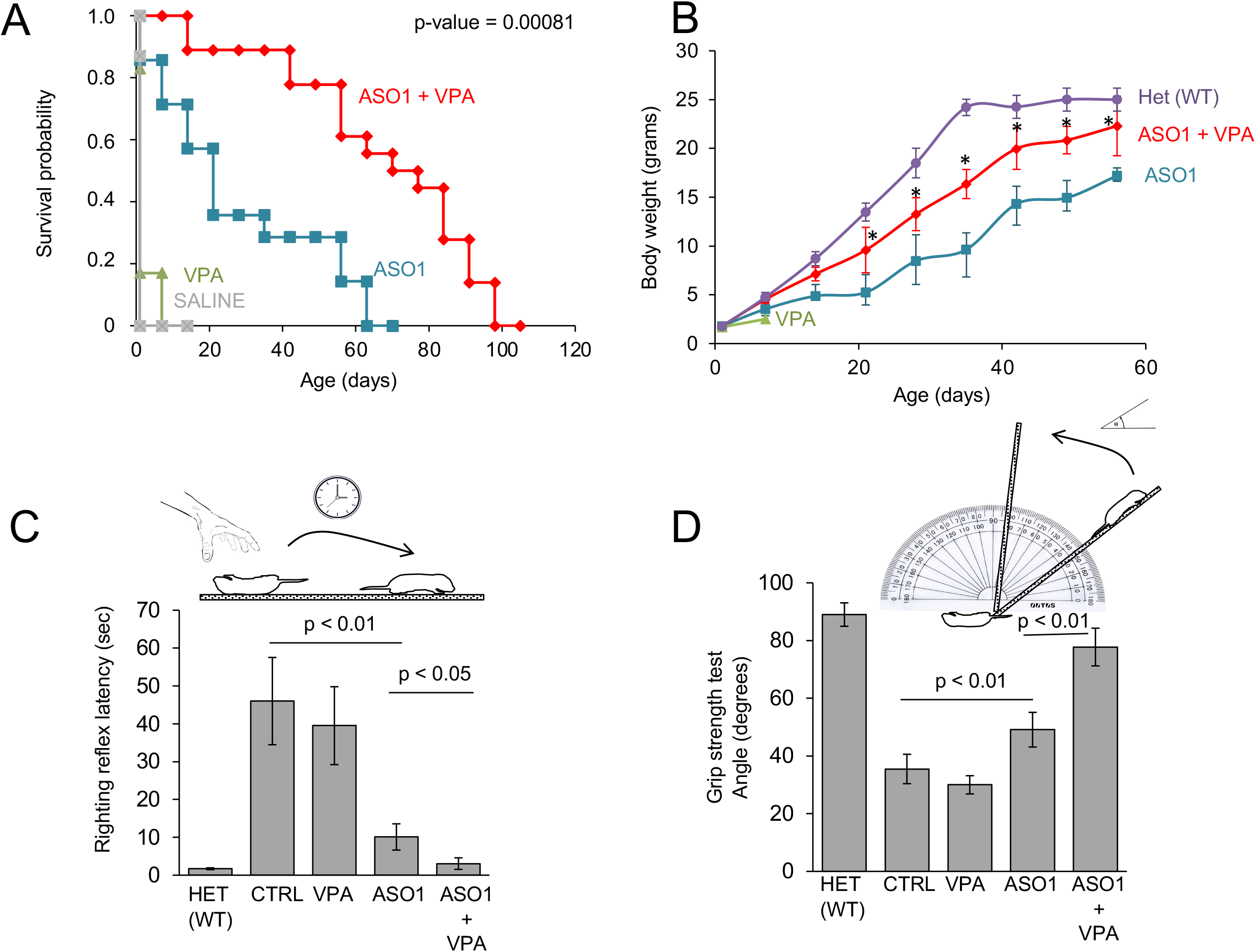
ASO1 and VPA have synergistic effects in SMA mice. Kaplan-Meier survival plot (A) and growth curves (B) of SMA mice, following subcutaneous administration at P0 and P1 of 16.8 μg ASO1 (n=18) or saline (n=14), one subcutaneous dose at P1 of 10 µg per g of body weight VPA (n=12), or both treatments together (n=20). ASO1-treated *Smn*^+ /-^ heterozygote littermates (n=15) served as controls. Righting reflex (C) and grip strength (D) tests of P7 SMA animals, treated as indicated in (A) and (B). In the righting-reflex test, we measured the time it takes a mouse to right itself when placed on its back on a flat surface. In the grip-strength test, we measured the angle at which the pups fall from a tablet with a rough surface to which they hold on by their forelimbs. Scramble (Ctrl) (n=12, 40 trials), VPA (n=13, 50 trials), ASO1 (n=11, 50 trials), ASO1 + VPA (n=17, 68 trials), and untreated heterozygotes (n=12, 36 trials). Statistical significance was analyzed by two-way repeated measures ANOVA. P < 0.05 was considered statistically significant; data are represented as mean + SD.

Body weight gain in the surviving mice was also greatly improved by VPA plus ASO1 injections, compared to ASO1 alone (Fig. 7B). We obtained similar results with a combined treatment of ASO1 and TSA (Suppl. Fig. 6A and B). Western blot analysis shows that the combined treatment greatly increased the levels of human SMN protein in liver, kidney, skeletal muscle, spinal cord, brain, and heart (Suppl. Fig. 6C-H). The largest effect on SMN expression was in the liver (Suppl. Fig. 6C), underscoring the proposed role of this organ in SMA pathogenesis (Hua et al., 2011).

Suppl. Fig. 7 illustrates conspicuous differences in size, posture, glassy eyes, and absence of grooming between the ASO1 alone and the ASO1+TSA-treated mice at 11, 14, 17 and 42 days after birth. Both the untreated and TSA-only-treated mice died around P7-P8.

We estimated the hazard ratio (HR) of the combined treatment versus a single drug treatment or no treatment at all. An HR greater than 1 suggests an increased risk, and an HR below 1 suggests a smaller risk. We obtained an HR = 0.19 for the combined treatment with VPA + ASO1 versus ASO1 alone, and an HR = 0.12 for the combined treatment with TSA + ASO1 versus ASO1 alone. Thus, far from being risky, these data demonstrate that the combined treatments are highly beneficial.

In addition, we performed two noninvasive neuromuscular function tests, appropriate for mouse pups. In the righting reflex test (Feather-Schussler et al., 2016), whereas wild-type mice righted immediately, untreated mutant mice took ∼45 seconds. This delay was not significantly changed by VPA treatment, was greatly reduced by ASO1 treatment (∼10 seconds), and was virtually eliminated by the combined treatment (Fig. 7C, Supplementary video 1). In the grip-strength test, animals treated with both ASO1 and VPA could hold on at much higher inclination angles (∼ 70º), compared with the untreated mutants (∼35º) or mice treated with VPA alone (∼30º) or limiting ASO1 alone (∼45º) (Fig. 7D, Supplementary video 2).

As in cultured cells (Fig. 3), ASO1 injection into SMA mice also promoted higher H3K9 dimethylation and RNAPII accumulation on the *SMN2* transgene in liver and brain (Suppl. Fig. 8). This result strongly suggests that the mechanism we elucidated in cultured cells also underlies the survival and behavioral effects observed in mice.

Recently, an additive effect on E7 inclusion of the HDAC inhibitor LBH589 and a nusinersen-like ASO was reported for SMA patient derived fibroblasts (Pagliarini et al., 2020). Similarly, the combination of VPA with a splice-switching phosphorodiamidate morpholino oligomer (PMO) was shown to promote more E7 inclusion than either agent alone (Farrelly-Rosch et al., 2017). However, neither study explored the underlying mechanism nor tested animal models.

### Therapeutical implications

Our results pave the way towards clinical assessments in patients. Although we expected VPA to have pleiotropic effects, our RNA-seq data of cells in culture indicated, in our conditions, that global effects on gene expression and splicing were minimal, which is consistent with our results in mice and with the fact that VPA is currently used to treat several neurological disorders (Chen et al., 2014). RNA-seq results reported here suggest that among HDAC inhibitors VPA is a good choice since it was reported that at molar concentrations 1000 smaller than VPA, other HDAC inhibitors like SAHA elicits expression changes in 10 times more genes than VPA, strongly affecting many signaling pathways (Lunke et al., 2021). In addition, although VPA has at best limited efficacy in SMA patients, it is safe and well tolerated (Elshafay et al., 2019). The lack of an effect *per se* in SMA mice (Figs. 7A and B) contrasting with the strong effect in improving ASO1’s therapeutic effects, suggests that an optimal VPA dose can be established that minimizes potential toxicities in patients. We speculate that systemic VPA administration will improve the efficacy of nusinersen, particularly in peripheral tissues. SMN is a ubiquitously expressed protein playing fundamental roles in splicing of all tissues. However, currently nusinersen is only administered via intrathecal injection to mainly reach the central nervous system. We foresee that, at low concentrations of nusinersen present in peripheral tissues, due to CSF clearance after intrathecal injection (Chiriboga et al., 2016), the combined systemic treatment with VPA or other HDAC inhibitors should increase SMN levels in the periphery, as shown in Suppl. Figs. 6C-H, and improve nusinersen performance in the periphery.

Finally we would like to underscore that our results support the relevance of the kinetic coupling between pre-mRNA processing and transcriptional elongation in a whole organism. Except for the alternative splicing/elongation response to light and dark in whole Arabidopsis plants (Godoy Herz et al., 2019) and non-physiological intervention with the slow polymerase mutant in mice (Maslon et al., 2019), most research supporting kinetic coupling was previously performed in cultured cells.

## Author contribution

L.E.M. performed most of the experiments with cultured cells and mice. Y.H.L handled the mice. G.D., J.S. and T.N. performed experiments with cells, N.J.P. supervised the design of genome-wide analyses and revised the manuscript and R.S-L. performed the bioinformatics analysis. A.R.Kr. and A.R.Ko. supervised the work. L.E.M., A.R.Kr., and A.R.Ko. wrote the manuscript.

## Acknowledgments

We thank Tassa Saldi and David Bentley for help with the re-analysis of their published RNA-seq data, Valeria Buggiano for technical assistance, and Julián Taranda for advice with the motor tests. This work was supported by a joint grant from Familias Atrofia Muscular Espinal (FAME, Argentina) and CureSMA (USA) and a grant from the Lounsbery Foundation (USA). A.R.Ko. acknowledges support from the Universidad de Buenos Aires (UBACYT 20020170100046BA), the Agencia Nacional de Promoción Científica y Tecnológica of Argentina (PICT-2019 862) and the CONICET (PUE 22920170100062CO). A.R.Kr. acknowledges support from NIH grant R37-GM42699 and the St. Giles Foundation. A.R.Ko. is a career investigator, and L.E.M. and J.S. received Ph.D. fellowships from the Consejo Nacional de Investigaciones Científicas y Técnicas of Argentina (CONICET). R.S-L. was funded by Fundação para Ciência e a Tecnologia, Portugal - Fellowship SFRH/BD/147906/2019. G.D. was supported by a Wellcome Trust Investigator Award to N.J.P (107928/Z/15/Z). A.R.Kr. is an inventor in nusinersen patents licensed by Cold Spring Harbor Laboratory to Ionis Pharmaceuticals, and sublicensed to Biogen, and is also a consultant to Biogen, which commercializes Spinraza.

## STAR METHODS

### Antisense oligonucleotide synthesis

ASO1 (5′-ATTCACTTTCATAATGCTGG-3′), a seven-mismatch control (5′-A<u>A</u>TCA<u>T</u>TT<u>G</u>C<u>T</u>T<u>C</u>AT<u>A</u>C<u>A</u>GG-3′) and ASO2 (5′-AAAGTATGTTTCTTCCACAC-3′) 2′-*O*-methoxyethyl-modified oligonucleotides with phosphorothioate backbone and all 5-methyl cytosines were purchased from IDT, and a 2′-OMe-modified phosphorothioate oligonucleotide (5′-AUUCACUUUCAUAAUGCUGG-3′)^1^ was purchased from TriLink. The oligonucleotides were dissolved in 0.9% w/v saline.

### RNAPII expression vectors and alternative splicing reporter minigene

The expression vectors for α-amanitin-resistant variants of the large subunit of human RNAPII (Rpb1) wild-type (WTres; pAT7Rpb1αAmr vector), and the hC4 mutant (pAT7Rpb1αAmrR749H) were previously described (de la Mata et al., 2003). An α-amanitin-sensitive Rpb1 expression vector (WTs) was used as a control. The pCI-*SMN2* (Addgene, Plasmid #72287) minigene vector for *SMN2* was described previously (Lorson et al., 1999).

### Cell culture and treatments

HEK293T, HeLa, and SMA type I homozygous and carrier fibroblasts 3813 and 3814 (Coriell Cell Repositories, Camden, New Jersey, United States) were grown in Dulbecco’s modified Eagle’s medium (DMEM) containing 4.5 g of glucose and 10% fetal bovine serum (Gibco) at 37 °C. Cells were plated at a density of 2×10^5^ cells per well in 12-well plates 24 hr before transfection. siRNA (25 nM), plasmid (500 ng), or ASO (25 nM) transfections were performed 24 hr after cells were plated, using 3 μl of Lipofectamine 2000 (Thermo Fisher Scientific) per well in 12-well plates. 24-48 h later, cells were treated with Trichostatin A (Sigma, T8552), Valproic Acid (Sigma, P4543), 5-Azacytidine (Sigma, A2385) or vehicle for the indicated time, and harvested for downstream procedures.

### RNA extraction and RT-PCR

Twenty milligrams of mouse tissue was pulverized in liquid N_2_ with mortar and pestle, or between 250,000 and 500,000 human cells, were homogenized with 1 mL of Trizol (Invitrogen). Total RNA was isolated according to the manufacturer’s directions. Two methods were used for RT-PCR reactions in different experiments: 1 One microgram of RNA was reverse-transcribed with M-MLV reverse transcriptase (Invitrogen) and oligo-dT primer, and the cDNA was amplified. Amplification and analysis of *SMN2* transcripts was performed as described1. 2: One microgram of total RNA was reversed transcribed using Superscript III (Thermo Fisher Scientific) reverse transcriptase and random primers. After PCR amplification using Gotaq (Promega) and *SMN2* primers surrounding E7, products were either loaded in a 6% acrylamide (37.5:1) TBE gel and stained with Ethidium Bromide for visualization or analysed by Tapestation 4150 using high sensitivity DNA D1000 screentapes (Agilent) for quantification.

### RNA-seq and library preparation

HEK293T RNA isolated as specified above were subjected to polyA RNA purification (NEBNext PolyA mRNA Magnetic Isolation Module, NEB). After purification, RNA spike-ins were added (SIRV-set2, Lexogene) to the pA+RNA. Then RNA was fragmented to 200 nt following the instructions of the NEBNext Ultra II Directional RNA library preparation kit to prepare PCR libraries. After libraries QC (Tapestation 4150, Agilent), NGS was performed on a Nova-seq platform (Novogene UK) to get 100 Mio PE reads per sample.

### POINT-seq

POINT-seq was performed as previously described (Sousa-Luís et al, 2020). In brief, HEK293T cells were either treated with VPA (10mM, 16hrs) or transfected with ASO1 (25nM, 48hrs). Chromatin-associated RNA was purified, followed by RNA Polymerase II immunoprecipitation to obtain Pol II-associated nascent transcripts. Isolated RNA were then mixed with spike-ins RNA (SIRV-set2, Lexogene) and NEBNext Ultra II Directional RNA library preparation kit was used to prepare PCR libraries. After libraries QC (Tapestation 4150, Agilent), NGS was performed on a Nova-seq platform (Novogene UK) to get 40-50 Mio PE reads per sample.

### Western blot

#### Mouse organs

Twenty milligrams of tissue was pulverized in liquid N_2_ and homogenized in 0.4 mL (liver, kidney, muscle, spinal cord, brain, and heart) of 1× protein sample buffer containing 2% (w/v) SDS, 10% (v/v) glycerol, 50 mM Tris-HCl (pH 6.8), and 0.1 M DTT. Protein samples were separated by 12% SDS-PAGE and electroblotted onto nitrocellulose membranes. The blots were probed with mAb anti-hSMN (BD Biosciences, 610646), or pAb anti-β-tubulin (Sigma), followed by secondary IRDye 800CW-conjugated goat anti-mouse or anti-rabbit antibody. Protein signals were detected with an Odyssey instrument (LI-COR Biosciences).

#### Cells in culture

Cells were lysed in 1X protein sample buffer (50mM Tris-HCl pH6.8, 2% SDS, 10% glycerol, 5% β-mercaptoethanol, 0.0025% Bromophenol blue). Protein samples were separated by 4-12% SDS-PAGE (NuPage, Life Technologies) and electroblotted onto PVDF membranes. The blots were probed with anti-H3K9me2 (Abcam, ab1220), anti-H3K9ac (Abcam, ab4441), anti-H3 (Abcam, ab12079), anti-Myc (Millipore, MABE282), anti-tubulin (Sigma-Aldrich, T5168), anti-RNase H1 (Proteintech, 15606-1-AP), anti-actin (Sigma-Aldrich, A2066), anti-G9A (CST, 3306S) or anti-Ago1 (CST, 5053S) antibodies.

### RNAi knockdown

Downregulation of hnRNP A1 and A2 was performed using ON-TARGET plus SMARTpool siRNA oligonucleotides (Dharmacon). siRNA oligos were delivered to cells following the manufacturer’s instructions, and allowed to act for 72 hr. Accell siRNA anti-human non targeting siRNA (Dharmacon, NC1567415) was used as a control.

siRNA oligos were delivered to cells following the manufacturer’s instructions, and allowed to act for 72 hr. siLuc (Sense: GAUUAUGUCCGGUUAUGUAUU Antisense: UACAUAACCGGACAUAAUCUU) was used as a control.

### Chromatin immunoprecipitation (xChIP) followed by q-PCR

Approximately 2 × 10^6^ HEK293T cells per sample were treated for 10 min in 1% (v/v) formaldehyde at room temperature to crosslink protein-DNA complexes. Crosslinking was stopped with glycine at a final concentration of 125 mM. Cells were washed twice with cold PBS and swelled on ice for 10 min in 25 mM HEPES pH 8, 1.5 mM MgCl_2_, 10 mM KCl, 0.1% NP-40, 1 mM DTT and 1× protease inhibitor cocktail set III (Calbiochem). Following Dounce homogenization, the nuclei were collected and resuspended in 1 ml sonication buffer (50 mM HEPES pH 8, 140 mM NaCl, 1 mM EDTA, 0.1% sodium deoxycholate, 0.1% SDS and 1× protease inhibitor cocktail). DNA was sonicated in an ultrasonic bath (Bioruptor Diagenode) to an average length of 200–500 bp. After addition of 1% (v/v) Triton X-100, samples were centrifuged at 15,000 ×g. Supernatants were immunoprecipitated O/N with 40 µl of pre-coated anti-IgG magnetic beads (Dynabeads M-280, Invitrogen) previously incubated with the antibody of interest for 6 hr at 4 ºC. The antibodies used were: rabbit anti-H3 (2 μg, Abcam ab1791), mouse anti-H3K9me2 (4 μg, Abcam ab1220), rabbit anti H3K9me3 (4 μg, Abcam ab8898), rabbit anti H3K9Ac (2 μg, Abcam ab4441), rabbit anti Rpb1 NTD (2 μg, Cell Signaling, D8L4Y), rabbit Phospho-Rpb1 CTD Ser2 (2 μg, Cell Signaling, E1Z3G), rabbit Phospho-Rpb1 CTD Ser5 (2 μg, Cell Signaling, D9N5I). Control immunoprecipitations were performed with rabbit IgG (1 μg, Abcam ab171870). Beads were washed sequentially for 5 min each in Low-salt (20 mM Tris-HCl pH 8, 150 mM NaCl, 2 mM EDTA, 1% Triton X-100, 0.1% SDS), High-Salt (20 mM Tris-HCl pH 8, 500 mM NaCl, 2 mM EDTA, 1% Triton X-100, 0.1% SDS) and LiCl buffer (10 mM Tris pH 8.0, 1 mM EDTA, 250 mM LiCl, 1% NP-40, 1% Na-deoxycholate) for 5 min at 4 ºC and then twice in TE 1× for 2 min at room temperature. Beads were eluted in 1% SDS and 100 mM NaHCO_3_ buffer for 15 min at 65 ºC and crosslinking was reversed for 6 hr after addition of NaCl to a final concentration of 200 mM. Chromatin was precipitated with ethanol overnight, treated with 20 μg proteinase K, and purified by phenol-chloroform extraction. Immunoprecipitated DNA (1.5 μl) and serial dilutions of the 10% input DNA (1:4, 1:20, 1:100, and 1:500) were analyzed by SYBR-Green real-time qPCR. The oligonucleotide sequences used are listed in Supplementary Table 1.

### Illumina data pre-processing

Quality control for raw Illumina short-reads was performed using the FastQC tool (https://www.bioinformatics.babraham.ac.uk/projects/fastqc/). Read adaptors were trimmed using TrimGalore (https://www.bioinformatics.babraham.ac.uk/projects/trim_galore/) in paired-end mode, removing reads with less than 10 nucleotides (nt) and/or low-quality ends (20 Phred scoref). The resulting reads were aligned against the reference human genome (GRCh38) using STAR software^2^, requiring a minimum alignment score (– outFilterScoreMin) of 10.

### Identification of expressed genes

Generated strand-specific pA+ RNA-seq data were used to identify expressed genes in the HEK293T cell line. Thus, *Kallisto*^3^ was used to map the reads against the human transcriptome (Ensembl v90), and TPM measurement for each transcription unit (TU) from the output was acquired. The transcript with highest TPM was selected for each gene. Genes having no transcript with TPM higher than 4 were discarded. Moreover, filtered TUs must have *protein-coding* tag as a biotype, which was extracted from Ensembl GTF file version 90. To better detect signal levels from POINT technology, overlapping TUs were excluded. To this end, an extra window of 10 Kb upstream and downstream of each TU was added. A final number of 2741 non-overlapping expressed genes was identified.

### Sashimi plots

For sashimi plots generation, ggsashimi v1.0.0 (https://doi.org/10.1371/journal.pcbi.1006360)^4^ was used on RNA-seq samples, with the following parameters: *-M 100 -s MATE2_SENSE -S plus*. The 3 replicates per condition were given as input. The number of read counts supporting each event was internally aggregated using the replicates mean. The following table shows the number of read counts per replicate, separated by a semicolon:

|  | Control | ASO1 | VPA | ASO1+VPA |
| --- | --- | --- | --- | --- |
| E6-E7 | 384; 930; 720 | 1002; 977; 994 | 1058; 1851;<br>1928 | 854; 2030; 1570 |
| E7-E8 | 452; 1018; 854 | 1327; 1053;<br>1040 | 1216; 1921;<br>2075 | 1008; 2313;<br>1818 |
| E6-E8 | 338; 1196; 636 | 370; 617; 361 | 1187; 970; 1172 | 339; 604; 441 |

### Meta-profiles

Previously identified expressed genes (N=2741 genes) were divided into 100 bins. For each bin, the overlapping number of fragment counts was measured with pysam^5^, and was then divided by the number of million reads mapped against the human genome, obtaining the *Fragment counts per Million Mapped Reads*.

Importantly, the 1% top and bottom extreme values per bin were removed, and the mean was measured excluding these values.

### POINT-seq window ratios

The promoter-proximal region was defined as a window from the TSS to 1500 bp downstream. The region from this point to the cleavage/polyadenylation site (pA site) was considered the gene body window. The gene set employed was the same as used in the meta-profiles (N=2741 genes). The number of fragment counts per window was obtained using pysam. Then, fragment counts were divided by the window length, to accommodate for differences in gene size. Thus, the ratio between promoter-proximal and gene-body regions was obtained by dividing the number of fragments in these respective windows, normalised to the window’s size. For the cases where no reads were found in the gene body, a value of zero was attributed. For the histogram, when there were no reads in the promoter-proximal region, a value of zero was obtained by adding 1 unit to the ratio and by measuring the log2.

The ratio between the regions upstream and downstream of the ASO1 target was obtained as described for promoter-proximal / gene-body regions. Here, the upstream window considered was *chr5:70951601-70951993* and the downstream was *chr5:70951993-70952250*.

### Identification of differentially expressed genes

Kallisto mapped RNA-seq reads against the human transcriptome to produce estimated gene-expression values, which were then gathered in a non-normalized count matrix.

Using it as input, significant differentially expressed genes were detected with the DESeq2 package (<u>10.1186/s13059-014-0550-8</u>). A cut-off of 10^−5^ for p-value and 1 for the absolute value of log2FoldChange was applied over DESeq2’s own two-sided statistical test results. Volcano plots were generated with the *EnhancedVolcano* v1.8.0 package (<u>10.18129/B9.bioc.EnhancedVolcano</u>).

### Differential alternatively spliced events

Genome-wide splicing variations induced by VPA treatment were investigated with rMATS v4.1.1^6^ Ensembl human v90 was given as annotations. The following parameters were also given to rMATS: -t paired --readLength 150 --libType fr-firststrand. The output was then stringently filtered by maser package (https://doi.org/doi:10.18129/B9.bioc.maser), such that an event was considered significant when the average number of reads per condition supporting the predominant class (inclusion or exclusion) was > 20, dPSI > 0.3 and FDR < 0.01. Scatter plot was obtained with the *dotplot* function of the maser package and represents the average PSI among the 3 replicates, per condition.

### Animals

All mouse protocols were in accordance with Cold Spring Harbor Laboratory’s Institutional Animal Care and Use Committee (IACUC) guidelines. The mild Hung mouse model (*Smn*^-/-^; *SMN2*^2TG /2TG^) was the strain FVB.Cg-*Smn1*^*tm1Hung*^ Tg(SMN2)2Hung/J, purchased from Jackson Laboratory (stock no. 005058). The severe SMA model (*Smn*^-/-^; *SMN2*^2TG /0^) was generated as previously described (Hua et al., 2011).

### Genotyping

For each animal, the genotype was verified by PCR reactions using tail-tip DNA. Primer sequences and PCR conditions were previously described^26^.

### Administration of oligonucleotides to hSMN2 transgenic mice

Oligonucleotide solutions in saline were injected subcutaneously into the upper back at P0 and P1 with a 5-mL syringe and 33-gauge custom removable needle (Hamilton) as described^2^. All drugs were injected subcutaneously before P3, contralateral to the oligonucleotide-injection site, with TSA (10 mg/kg) or VPA (10 mg/kg) or vehicle. Mouse tissues and organs, including liver, thigh muscles, kidney, and spinal cord, were snap-frozen in liquid N_2_ and stored at −70 °C.

Power calculations were done according to standard requirements for animal-protocol approval by CSHL’s IACUC. Based on anticipated effect sizes, we aimed to have 12-15 pups per group for the survival analysis. Because the crosses to obtain SMA mice yield ∼50% of non-SMA heterozygote mice, we needed a total of 120 pups to obtain ∼60 SMA mice (15 pups x 4 treatments) for each HDAC inhibitor treatment with its own controls. To obtain this number of mice, a total of 60 breeding cages were used.

The number of 12-15 pups per group was decided following the conditions of Hua et al., 2011. This number was chosen in order to test the effect of VPA/TSA on ASO1 therapy with inferential statistics (i.e., using p-values) and at the same time to avoid sacrificing an excessive number of mice, according to https://research.usu.edu//irb/wp-content/uploads/sites/12/2015/08/A_Researchers_Guide_to_Power_Analysis_USU. pdf and to the NIH Guide for the Care and Use of Laboratory Animals (2011).

## Notes

### Competing Interest Statement

The authors have declared no competing interest.

