## Supplementary material for "Towards a combined therapy for spinal muscular atrophy based on opposing effects of an antisense oligonucleotide on chromatin and splicing": SUPPL. FIGURES + TEXT

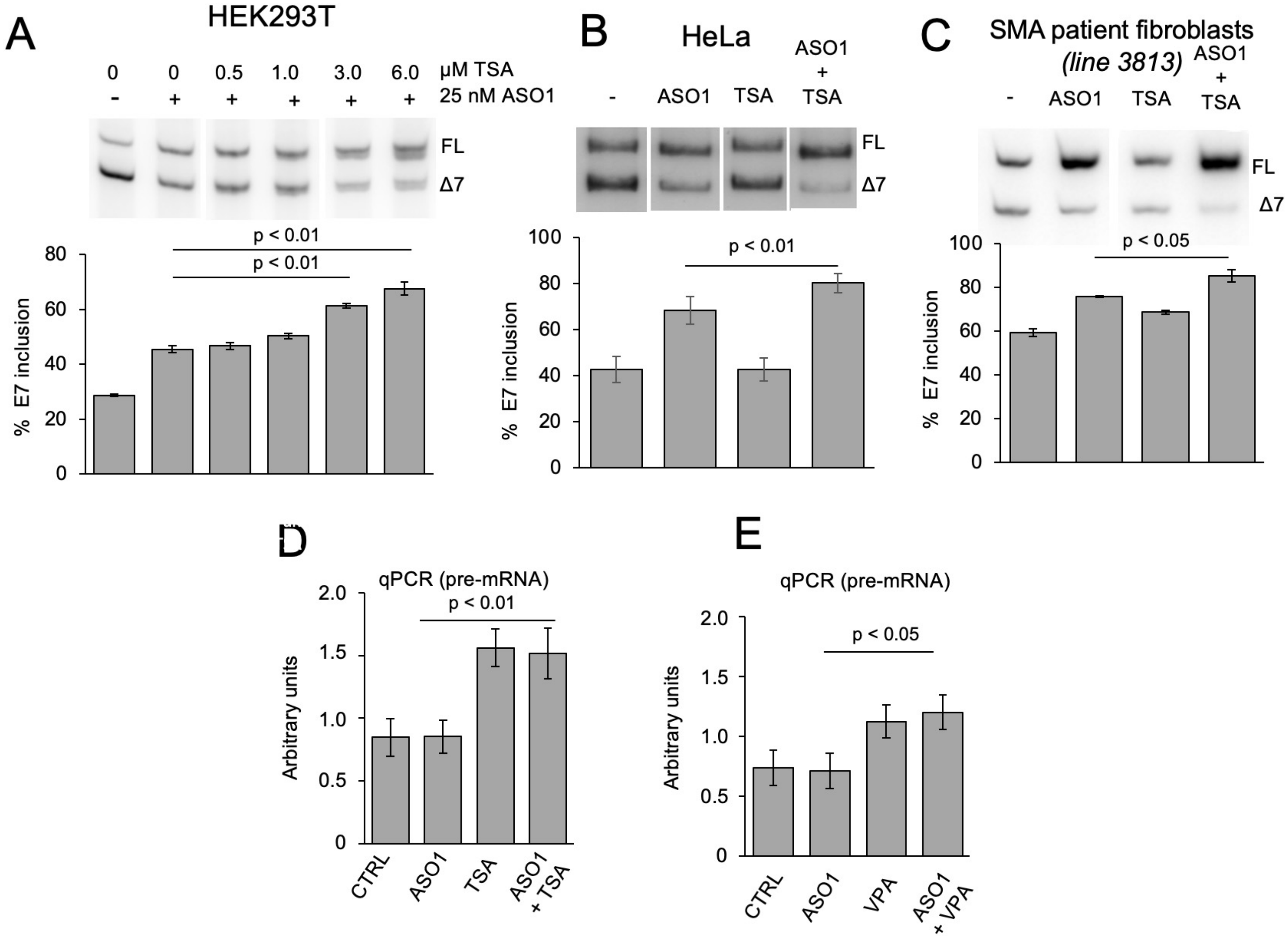


**Supplementary Figure 1. Chromatin relaxation potentiates the ASO effect on E7 inclusion.** Combined effects on endogenous *SMN2* E7 alternative splicing in transfected (A) HEK293T cells treated with 25 nM ASO1 and increasing doses of TSA for 24 hr, (B) HeLa cells treated with 10 nM ASO1 and 3 $\mu$M TSA for 24 hr., and (C) SMA patient fibroblasts (3813) treated with 10 nM ASO1 and 3 $\mu$M TSA for 24 hr. Bars display means ± SD of percentage of the radioactivity in the FL band over the sum of radioactivity in the FL and DE7 bands of at least three independent transfection experiments. (Dand E). Transcriptional activation was assessed by RT-qPCR. Gray bars indicate mean transcriptional activation ± SD ratios between precursors of *SMN2* and *GAPDH* as the control gene. Statistical significance was analyzed by two-tailed Student's t-tests. P<0.05 was considered statistically significant.

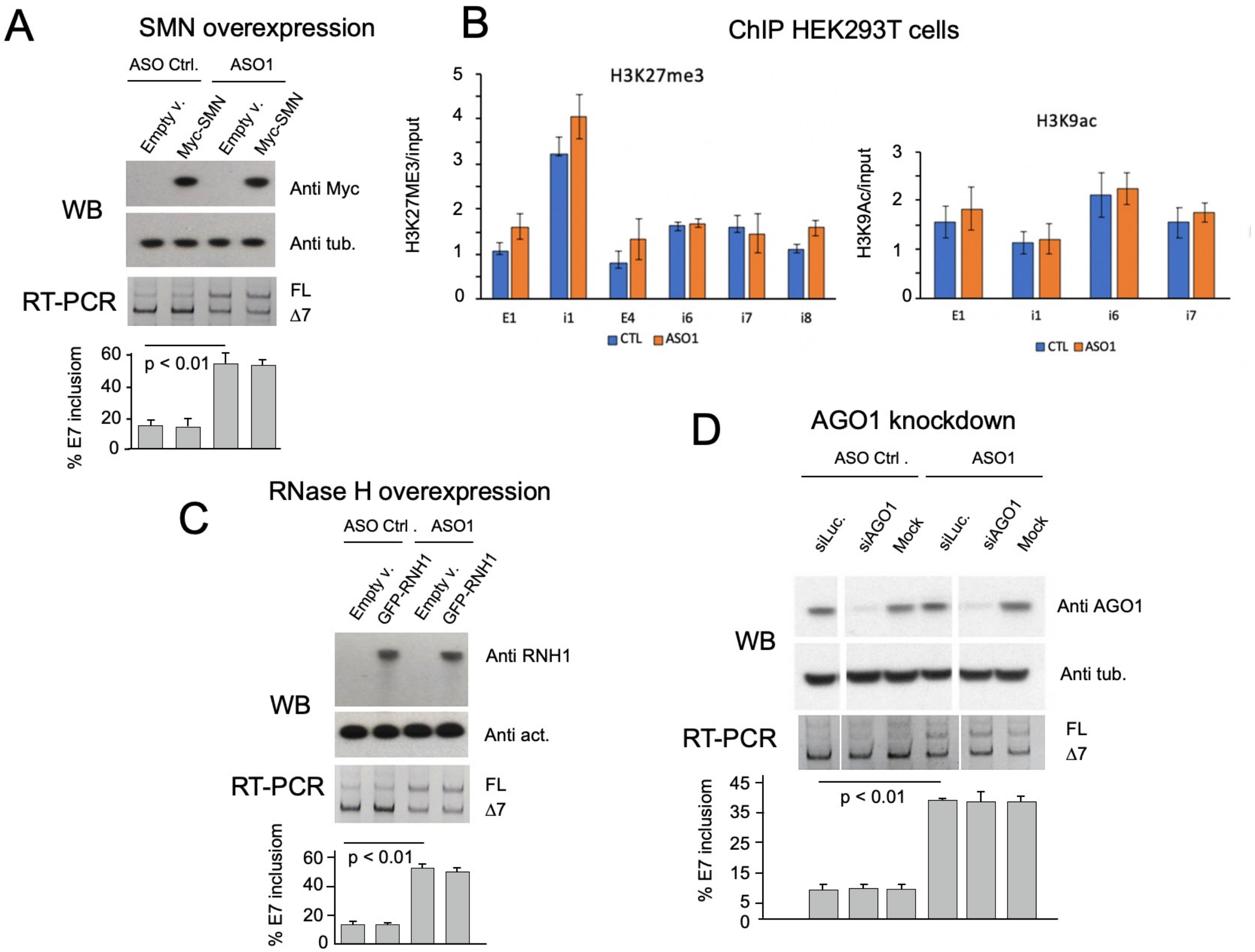


**Supplementary Figure 2. Control experiments for the opposing roles of ASO1 on chromatin and splicing.** (A) Overexpression of the SMN protein does not alter the ASO1 effect on E7 splicing. Top: control Western blot for efficient expression of the c-Myc-SMN fusion protein encoded by plasmid pcDNA3.1+SMN1mycHIS (Addgene, cat.# 71687) using anti-tubulin (tub.) as loading control. Bottom: *SMN2* E7 RT-PCR of cells co-transfected with ASO1 and plasmid. (B) Transfection of HEK293T cells with ASO1 does not increase H3K27 trimethylation or H3K9 acetylation over the *SMN2* gene. Distribution of these histone marks was assessed by ChIP-qPCR, with amplicons mapping near the numbered exons (E) and introns (i). Three independent immunoprecipitation replicates were conducted per experiment. Data are represented as mean ± S.D. ($n = 5$; in all cases p was much higher than 0.05, two-tailed Student's t test). (C) Overexpression of the RNase H enzyme does not affect the ASO1 effect on E7 splicing. Top: control Western blot for efficient expression of the GFP-RNase H fusion protein encoded by plasmid pEGFP-RNASEH1 (Addgene, cat.# 108699), using anti-actin (act.) as loading control. Bottom: *SMN2* E7 RT-PCR of cells co-transfected with ASO1 and the RNase H plasmid. (D) siRNA-mediated AGO1 knockdown does not alter the ASO1 effect on E7 splicing. Top: control Western blot for efficient AGO1 knockdown using anti-tubulin (tub.) as loading control. Bottom: *SMN2* E7 RT-PCR of cells co-transfected with ASO1 and the AGO1 siRNA. RT-PCR conditions for panels **a**, **c** and **d** were as in main Figure 1A.

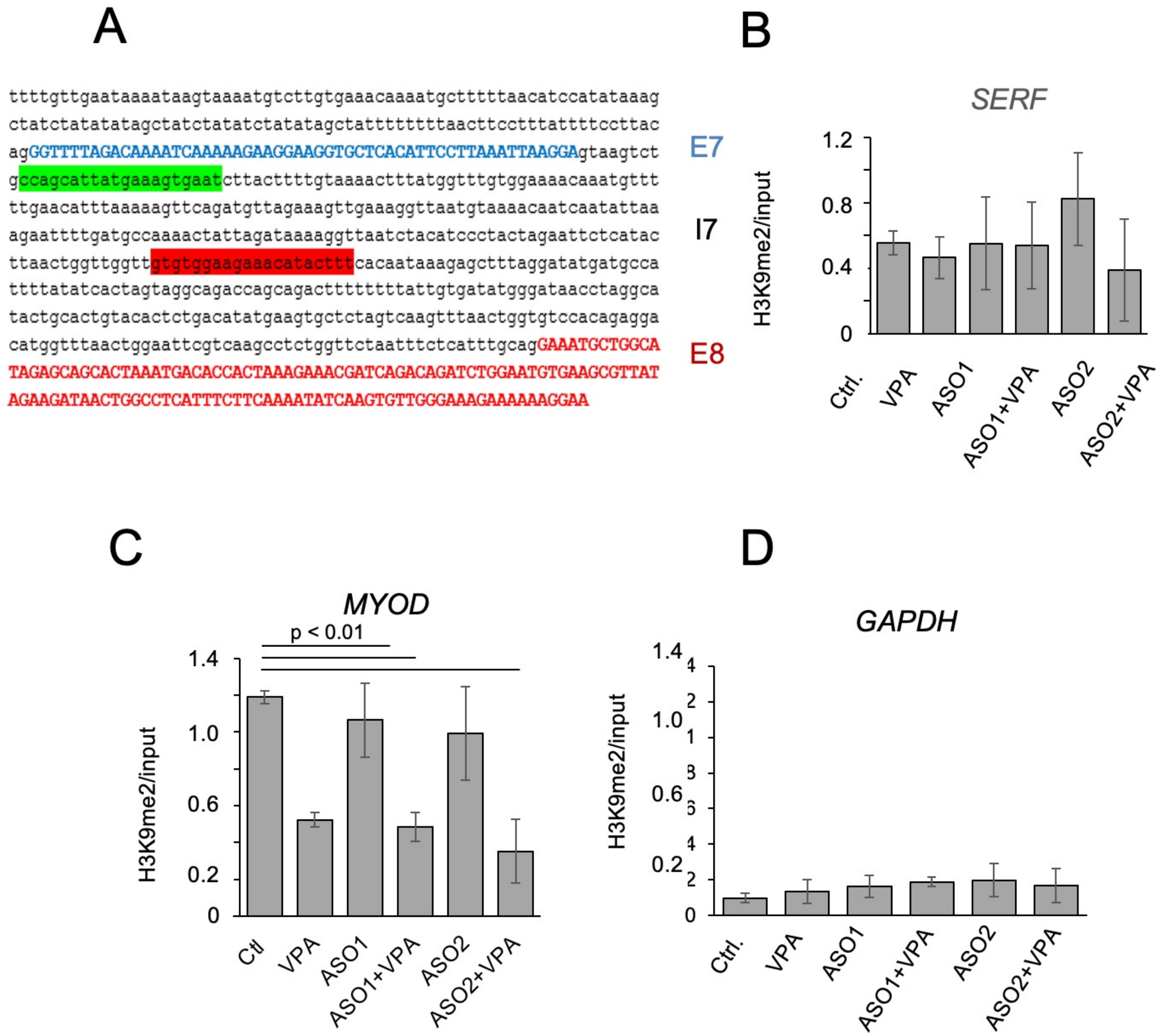


**Supplementary Figure 3. Antisense oligonucleotides and their effects on chromatin.** (A) Exert of the *SMN2* gene sequence comprising the E7 alternative exon. Blue uppercase, exon 7 nucleotide sequence. Lowercase, intronic sequences. Red uppercase, exon 8 nucleotide sequence. Green highlight, sequence corresponding to the binding site for ASO1 and hnRNPA1/A2 i the encoded pre-mRNA. Red highlight, idem for the binding site for ASO2. (B-D) Levels of H3K9me2 deposition along (B) Small EDRK-rich factor 1A (*SERF1A*, upstream of *SMN2*), (C) Myoblast Determination Protein (*MYOD*), and (D) Glyceraldehyde 3-phosphate dehydrogenase (*GAPDH*), assessed by ChIP-qPCR in HEK293T cells. Data represented as mean ± S.D. ($n = 4$, $^*p < 0.05$, two-tailed Student's t test).

A

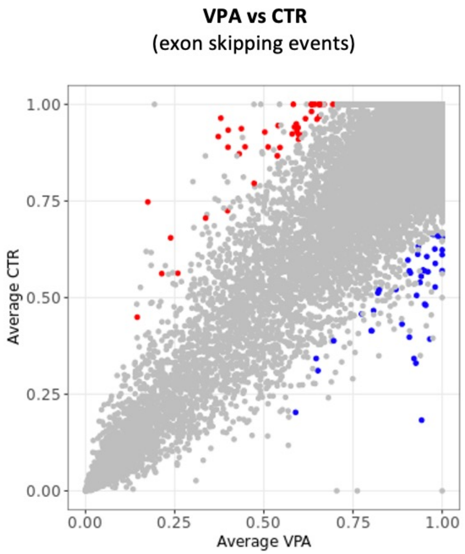


B

| AS mode | # Total ASE | # Dif. ASE | % Dif. ASE |
|---|---|---|---|
| alternative 3' ss | 4494 | 4 | 0.0890% |
| alternative 5' ss | 3018 | 9 | 0.2982% |
| cassette exon | 58668 | 33 | 0.0562% |
| intron retention | 3239 | 9 | 0.278% |
| mutually exclusive exons | 10130 | 2 | 0.0197% |
| TOTAL | 79549 | 48 | 0.0603% |

C

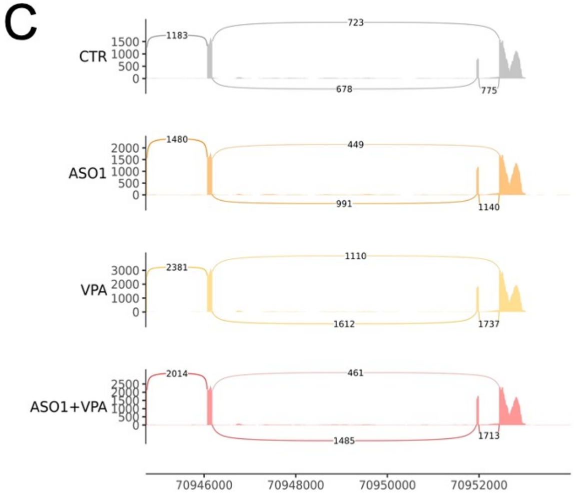


D

E7 inclusion analyzed through E6-E7 junction reads

| Treatment | E6-E8 junction reads (E7 skipping) | E6-E7 junction reads (E7 inclusion) | % E7 inclusion | Corrected % E7 inclusion | Corrected E7+/E7- ratio |
|---|---|---|---|---|---|
| Control | 723 | 678 | 48.3 | 3.4 | 0.04 |
| ASO1 | 449 | 991 | 68.8 | 37.6 | 0.6 |
| VPA | 1110 | 1612 | 59.2 | 18.4 | 0.2 |
| ASO1+ VPA | 461 | 1485 | 76,3 | 52.6 | 1.1 |

**Supplementary Figure 4. Global effects of VPA on alternative splicing revealed by RNA-seq.** (A) Scatter-plot showing PSI (Percentage Spliced-In) of exon skipping events detected by rMATS, for RNA-seq samples. Red dots are differentially spliced events for which the inclusion is higher in the control cells, and blue dots represent events for which the inclusion is higher in VPA-treated cells. Grey dots represent not significant cases. Cases were only deemed significant if the number of supportive reads > 20, dPSI > 0.3 and FDR < 0.01. (B) Table showing the number of alternative splicing events (ASEs) of different modes, whose patterns are significantly affected by VPA. Total ASE are all alternative splicing events with more than 20 reads. Differential ASE are the ones for which supportive reads were > 20, dPSI > 0.3 and FDR < 0.01. ss: splice site. (C) Sashimi plot showing the average number of reads supporting each splice junction of the *SMN1/2* merged genes, based on the aligned RNA-seq data, upon treatments of HEK293T cells with ASO1, VPA or both together. (D) Raw and corrected quantification of the levels of E7 inclusion as assessed by analysis of the E6-E7 junction reads. Inclusion levels are expressed as both percentage of E7 inclusion and E7+/ E7- (i.e. FL/Δ7) ratios.

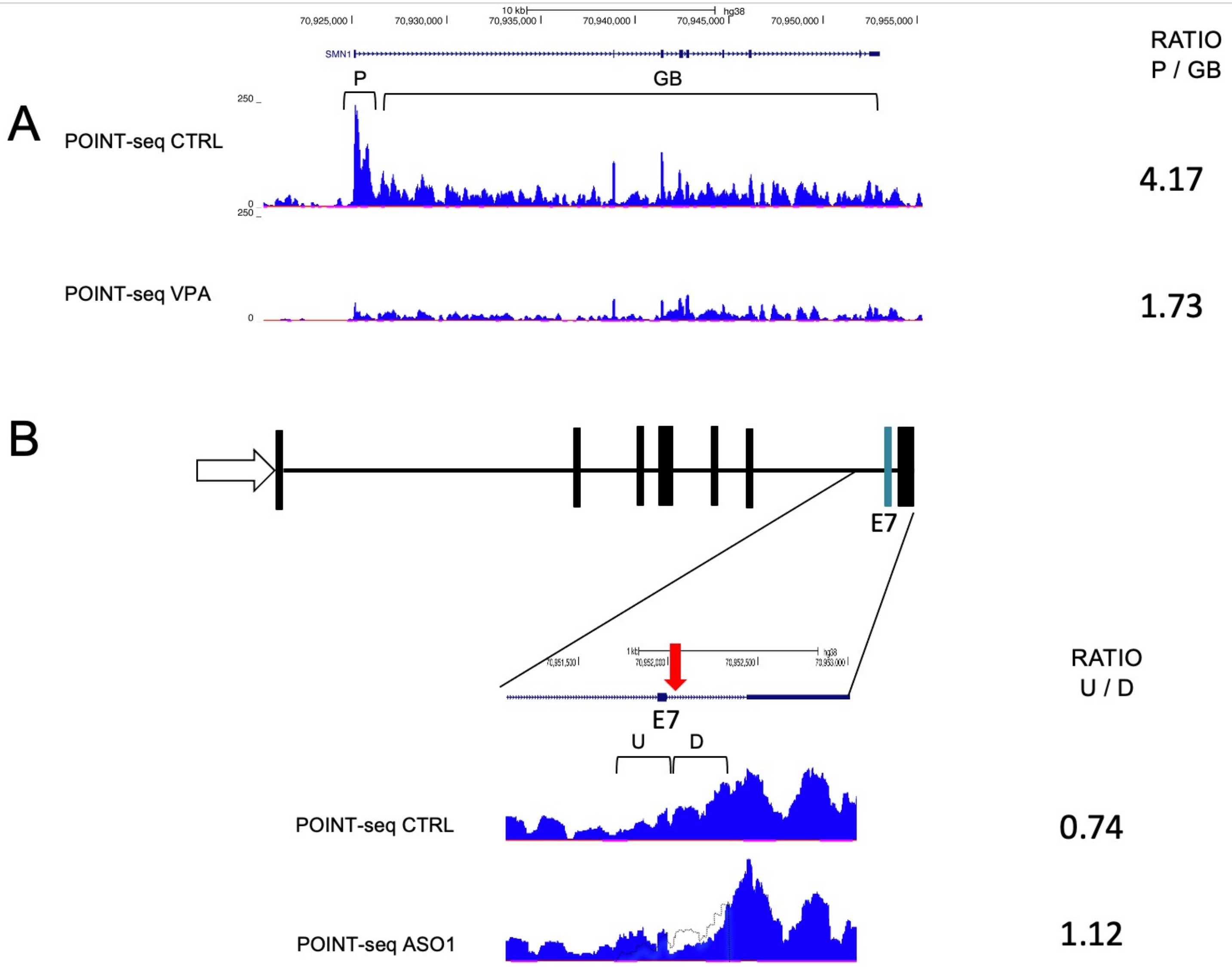


**Supplementary Figure 5. POINT-seq analysis on the SMN1/2 merged genes.** (A) POINT-seq signal normalized to the library size on the *SMN1/2* merged genes of HEK293T cells treated with VPA. Ratios between the accumulated reads at the promoter region (P) and gene body (GB) are shown on the right. (B) POINT-seq signal at a zoom-in of the target region of ASO1, displaying the impact of ASO1-binding on Pol II progression. The top track is the control sample, and the bottom corresponds to ASO1-treated cells. Read density ratios of the upstream (U) over the downstream (D) regions are shown on the right side of the corresponding track. The control profile is superimposed as a dotted line on the ASO1 profile.

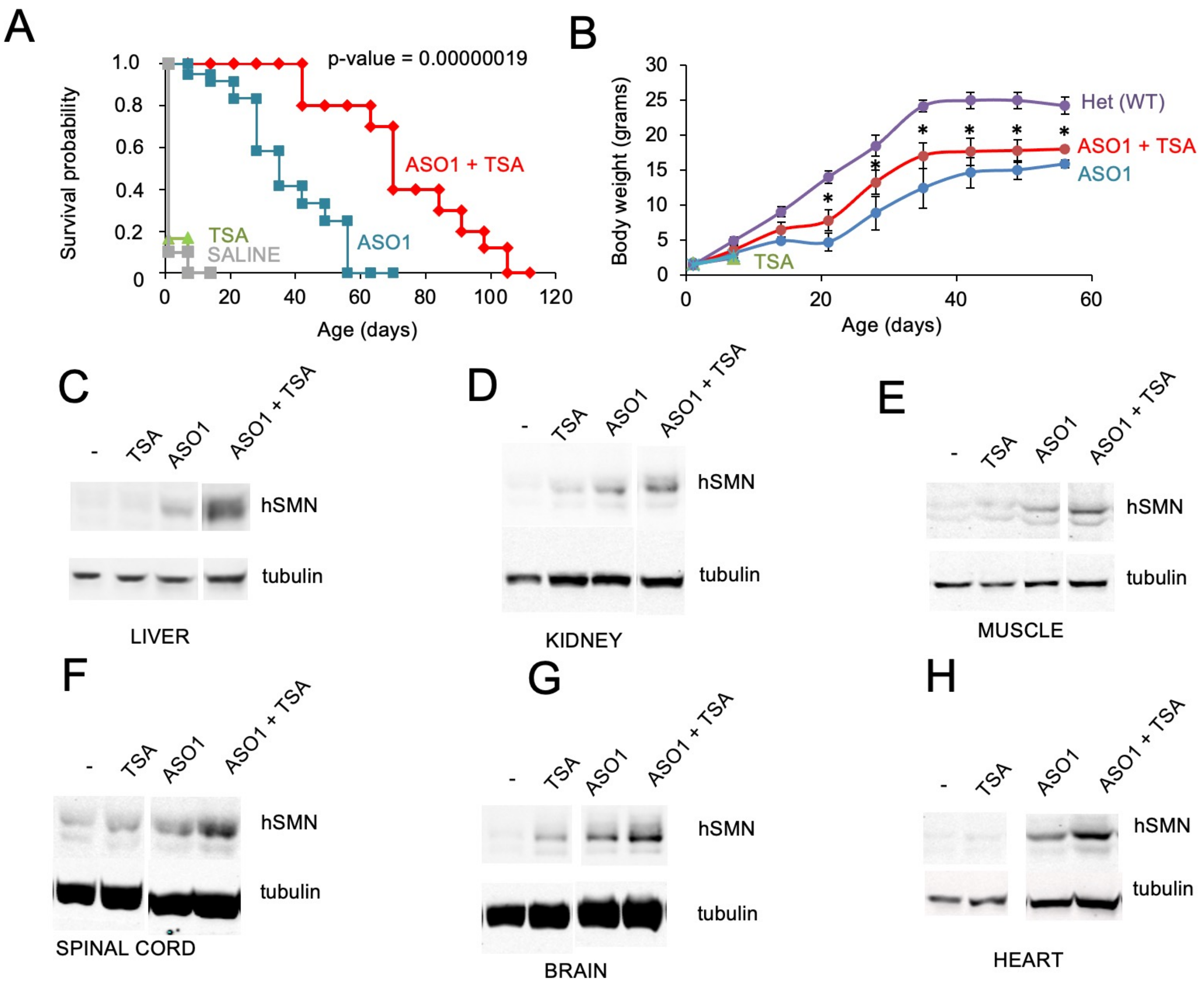


**Supplementary Figure 6. ASO1 and TSA combined treatment in mice.** Kaplan-Meier survival plot (A) and growth curves (B) of SMA mice after subcutaneous administration at P0 and P1 of 16.8 μg ASO1 (n=20) or vehicle (n=12), one subcutaneous dose of 10 μg per g of body weight TSA (n=15) at P2, or both treatments together (n=24). ASO1-treated heterozygotes (n=18) served as controls. Statistical significance was analyzed by two-way repeated measures ANOVA. $P < 0.05$ was considered statistically significant; data are represented as mean + SD. **(C-H)** Western-blot analysis of P7 tissues from liver (C), kidney (D), muscle (E), spinal cord (F), brain (G) and heart (H) from SMA mice, using anti-hSMN (BD Biosciences, upper panel) and anti-β-tubulin (Sigma, lower panel) antibodies.

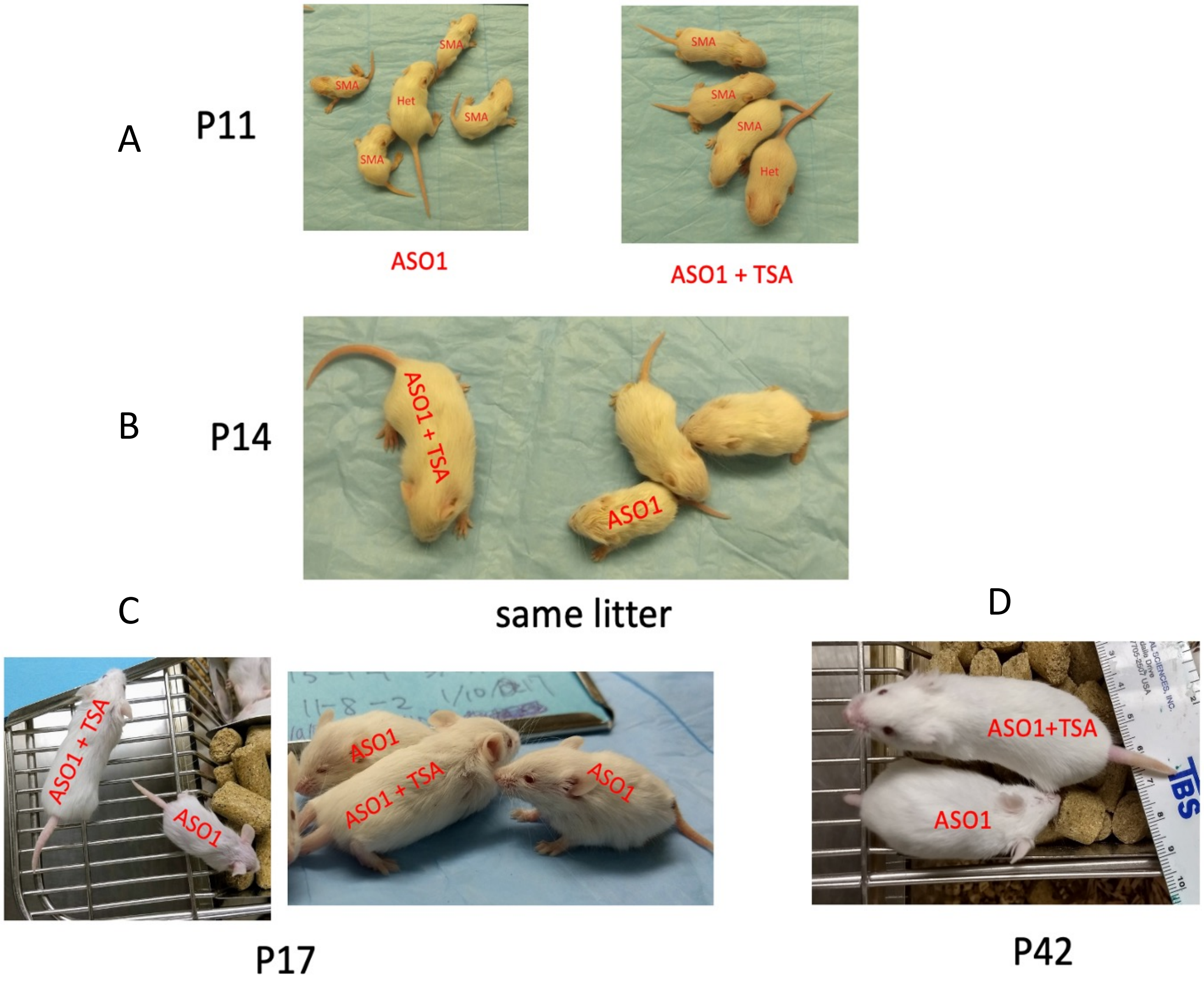


**Supplementary Figure 7.** Phenotypes of mice treated with subotimal doses of ASO1 alone or in combination with TSA. (A) Comparison of body sizes at P11 of mice heterozygous (Het) and homozigous (SMA) for the mutated mouse *SMN* gene inhected with ASO1 (left) or ASO1 + TSA (right). (B-D) Phenotypes of SMA mice at P14 (B), P17 (C) and P42 (D) treated with ASO1 or ASO1 + TSA.

SMA mice P7

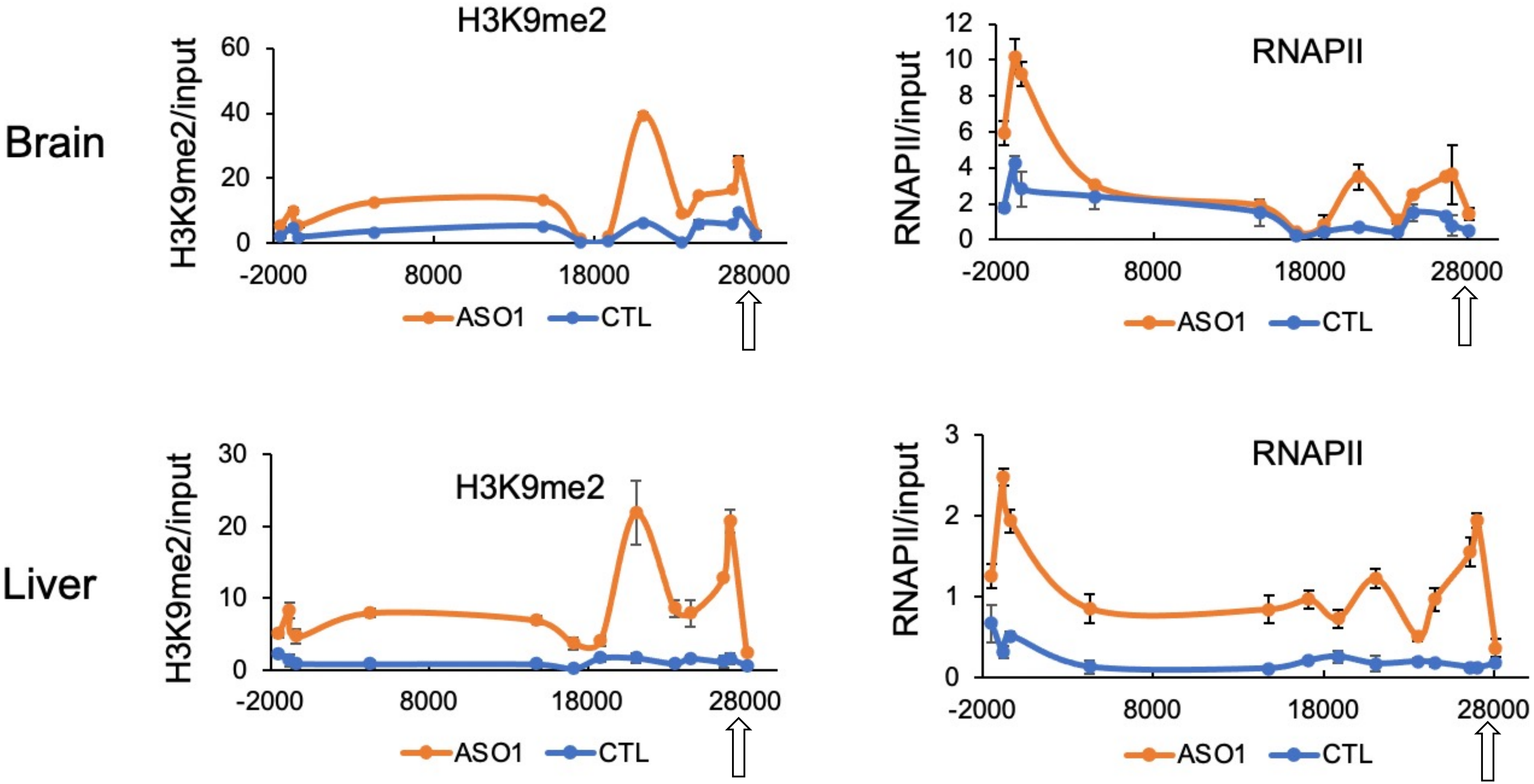


**Supplementary Figure 8. SMA mice injected with ASO1 show increased H3K9 dimethylation and RNAPII roadblocks along the *SMN2* gene in brain and liver.** H3K9me2 (left) and total RNAPII (right) distribution along the human *SMN2* transgene, assessed by ChIP-qPCR, in brain (top) and liver (bottom) of P7 SMA mice injected with ASO1 as indicated in Methods. Vertical arrows indicate the approximate location of the target site for ASO1 on the pre-mRNA. Three independent immunoprecipitation replicates were conducted per experiment. Data are represented as mean ± S.D. ($n = 3$, $^*p < 0.05$, two-tailed Student's t test).
